# Pan-cancer analysis reveals that tumor microbiomes generate specific epitopes through transcriptional reprogramming

**DOI:** 10.1101/2025.10.22.683275

**Authors:** Jinman Fang, Yuxin Zhang, Wei Wang, Xiao Zuo, Tao Chen, Kecheng Li, Huiya Yang, Tianyang Zhu, Yuanliang Zhang, Dandan Wei, Jianfeng Zhang, Jialong Cui, Zhen Tang, Shouwu Zhang, Lirong Wang, Wei Wang, Zhilei Li, Dushan Ding, Ruixia Sun, Dandan Zhu, Yin He, Yanfei Zhou, Chenkun Ye, Dandan Chen, Xueting Lang, Long Jiang, Qian Zhao, Lu Zhang, Hongzhi Wang, Yuanwei Zhang

## Abstract

The therapeutic promise of tumor microbial peptides faces a critical safety paradox: tumors and peritumor tissues harbor compositionally similar microbiomes, yet safe immunotherapy demands tumor-specific targeting. We performed comprehensive transcriptomic analysis of microbial communities from 1,868 tumor and peritumor samples across 11 cancer types. Although 88.5% of microbial species are present in both tissue types, tumor and peritumor tissues exhibit profound transcriptional divergence: only 34.1% of microbial genes are expressed in both tissue types, and merely 18.2% of predicted HLA-binding peptides overlap. This transcriptional divergence represents microbial transcriptional reprogramming (MTR) in tumor tissues, through which microbes activate distinct genomic regions while adapting to tumor microenvironmental pressures, thereby generating tumor-specific peptide repertoires. Using Microbial Transcriptional Reprogramming Index (MiTRI) to quantify MTR, we found that higher MiTRI values correlate with expanded immunogenic peptide repertoires. Mass spectrometry-based immunopeptidomics confirmed that 25% of naturally presented microbial peptides originate from MTR regions. Crucially, in microsatellite-stable colorectal cancer, neo-adjuvant therapy responders exhibited significantly higher MiTRI values. T cell receptors (TCRs) recognized MTR-derived epitopes at 2.10-fold higher rates than genome-wide epitopes, accounting for 79.1% of clonally expanded TCRs. These findings establish MTR as the principle that ensures the tumor-specificity and safety of micorbiome-directed immunotherapy.

## Introduction

Targeting microbial peptides has emerged as a promising immunotherapy strategy for cancer treatment. Recent studies demonstrated that tumor-resident bacteria present peptides on major histocompatibility complex (MHC) molecules that can elicit anti-tumor immune responses^1–10^, and mouse models showed that killing these bacteria with antibiotics can expose microbial epitopes driving therapeutic immunity^6^. These findings have catalyzed clinical translation, with microbiome-derived molecular-mimicry peptide vaccines now entering early-phase trials ^11,12^. However, a fundamental challenge threatens the safe clinical deployment of this approach: tumors and peritumor tissues harbor remarkably similar microbial communities ^13–15^, yet the principle ensuring safe, tumor-specific targeting of microbial peptides remains unknown..

This knowledge gap stems from a critical disconnect in perspective. When studying microbes as tumor-promoting factors, the focus has traditionally been on their enrichment in tumor tissues^1–10^, irrespective of whether such microbial presence is specific or similarly present in peritumor tissues. However, when shifting the perspective to tumor peptides, particularly microbial peptides considered for their anti-tumor potential, the emphasis must pivot towards specificity. This shift is critical to ensuring the safety and efficacy of clinical applications, such as in immunotherapeutic strategies. Currently, there remains a notable scarcity of large-scale, pan-cancer studies systematically investigating the specificity of microbial peptides ^16^. More importantly, the field still lacks a coherent theoretical framework to explain the specificity of microbial-derived peptides, unlike the well-established paradigm of somatic mutations generating tumor-specific neoantigens.

Here, we performed pan-cancer transcriptomic analysis of 1,868 tumor and peritumor samples across 11 cancer types. We discovered that while 88.5% of microbial species residented in both tumor and peritumor tissues, only 34.1% of microbial genes are expressed in both tissue types. We hypothesized that tumor microenvironments drive microbial transcriptional reprogramming (MTR) of resident microbes, generating tumor-specific peptide repertoires distinct from those in peritumor tissues. This concept is grounded in the well-established principle that microorganisms adapt to environmental niches through selective gene activation ^17,18^, observed in contexts ranging from bacterial stress responses to host-microbiome co-evolution ^19,20^.

To quantify the MTR degree of microbes, we developed the microbial transcriptional reprogramming index (MiTRI) and linked it to immunogenic output. Mass spectrometry-based immunopeptidomics validated the natural presentation of predicted microbial peptides, with a 25% enrichment for those derived from MTR regions. Crucially,in microsatellite-stable (MSS) colorectal cancer (CRC), a setting with scarce conventional neoantigens, neo-adjuvent therapy responders exhibited significantly higher MiTRI, and T cell receptors (TCRs) recognized MTR-derived epitopes at a 2.10-fold higher rate than genome-wide epitopes. Our work thus establishes MTR as a unifying principle that ensures the specificity and safety of microbiome-directed immunotherapy.

## Results

### Compositionally similar microbiomes shapes a tumor-specific microbial peptide landscape

To compare microbial peptide landscapes between tumor and peritumor tissues, we performed comprehensive transcriptomic analysis of 1,169 tumor and 699 peritumor samples across 11 cancer types (**Methods**; **Extended Data Table S1**), with a curated tumor microbial genomic reference dataset (**Methods**; **Extended Data Fig. S1** and **Extended Data Table S2**) ^13,21^. We identified 2,775 microbial species with 25 million predicted microbial human leukocyte antigen (HLA)-binding peptides (**Methods**), using a computational pipeline for identifying microbes and infering microbial peptides from whole-transcriptome sequencing (WTS) ^22^.

As expected from previous studies ^13–15^, we observed that tumor and peritumor tissues harbored compositionally similar microbial communities, with 93.7% of genus and 88.5% of species resident in both tissue types (**Methods; Fig. 1A-B**). These proportions vary from 55.6% in cervical cancer (CESC) to 94.2% in CRC for genus (**Extended Data Fig. S2**), and from 34% in CESC to 90.5% in CRC for species (**Extended Data Fig. S3**). However, beneath this compositional similarity, microbial gene expression profiles differed fundamentally (**Methods**). Across all 11 cancer types, only 34.1% of microbial genes were expressed in both tumor and peritumor tissues (**Methods; Fig. 1C**), with proportions ranging from 3.5% in bladder cancer (BLCA) to 44.5% in renal cell carcinoma (RCC) (**Extended Data Fig. S4**). Among the 2,455 species resident in tumor and peritumor tissues, only 36.7% of expressed genes were expressed in both tissue types (**Fig. 1D**), with proportions varying from 5.9% in PAAD to 45.5% in CRC (**Extended Data Fig. S5**). We observed this environment-driven transcriptional divergence reshaped transcriptional outputs, where Gene Ontology (GO) enrichment analysis on specifically expressed genes revealed 491 terms significantly enriched in tumor tissues and 244 in peritumor tissues, with only five shared terms between them (FDR-BH < 0.05; **Methods**; **Extended Data Table S3**).

**Fig. 1.**
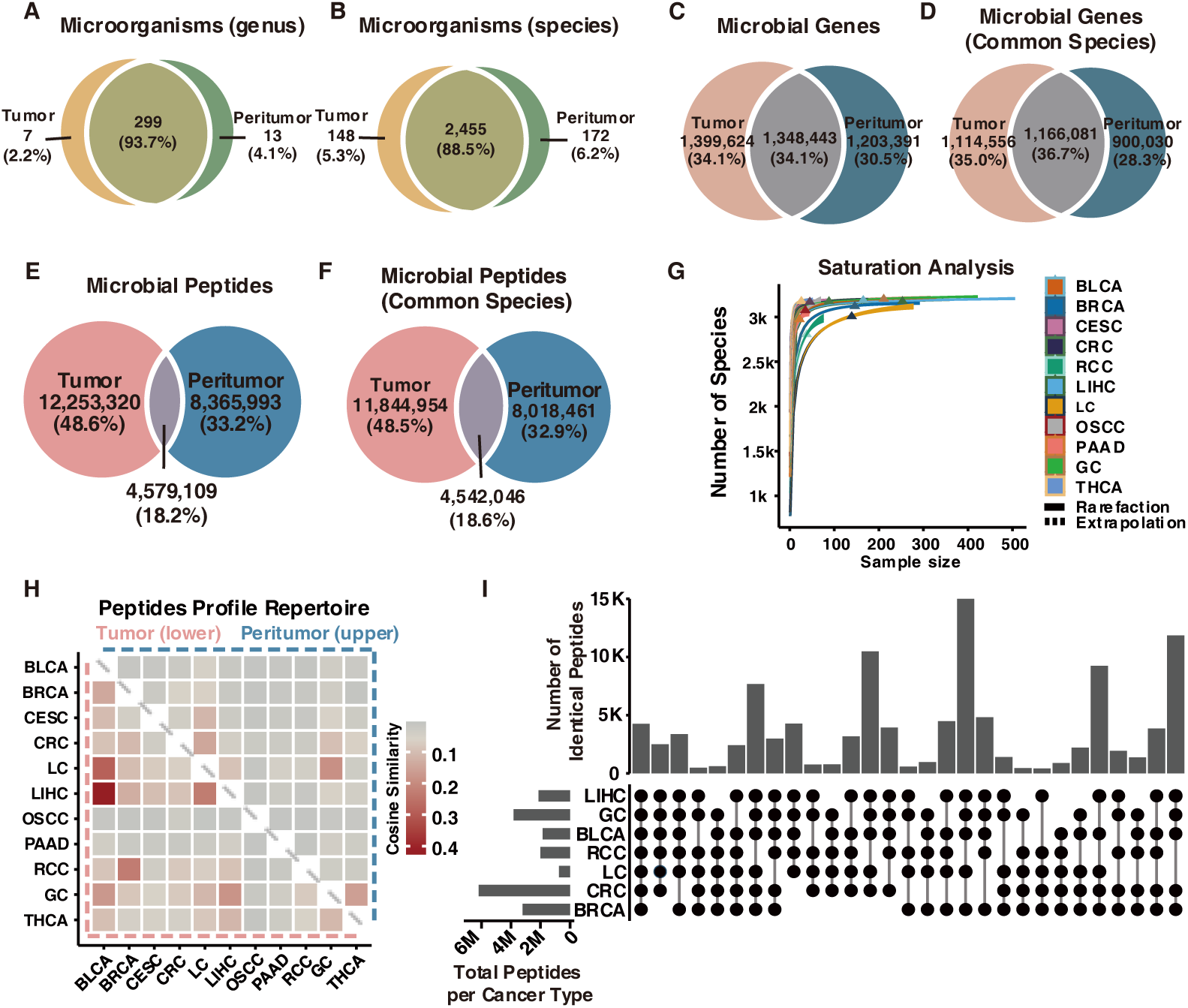
Microbiome composition in tumor and peritumor tissues across cancer types. **(A-B)** Venn diagrams show the microbial composition at the genus (A) and species (B) levels between tumor and peritumor tissues across 11 cancer types. 93.7% of genera and 88.5% of species are present in both tumor and peritumor tissues across 11 cancer types (Microbes were considered “present” when detected in ≥50% of samples from each tissue type.) **(C)** Venn diagram shows 34.1% of microbial genes expressed in both tumor and peritumor tissue samples. **(D)** Venn diagram shows the 36.7% of microbial genes expressed in both tumor and peritumor tissue samples from common microbial species identified in panel B (“common species” refers to microbial species detected in both tumor and peritumor tissues). **(E)** Venn diagram shows the 10.5% of shared predicted HLA-binding peptides between tumor and peritumor tissue samples. **(F)**Venn diagram shows the 18.6% of predicted HLA-binding peptides from common microbial species identified in panel B. **(G)** Rarefaction curves show the observed species saturation across 11 cancer types. **(H)** Heatmap of the cosine similarity matrix comparing predicted microbial HLA-binding peptide repertoires in tumor and peritumor tissue samples. Tumor samples show significantly higher intra-group similarity (maximum = 0.42) than peritumor tissues (maximum = 0.19; *p* = 0.0015, one-tailed Mann–Whitney test). **(I)** UpSet plot shows overlaps of predicted microbial HLA-binding peptides exclusively detected in tumor samples. Abbreviations: bladder cancer, BLCA; breast cancer, BRCA; cervical cancer, CESC; colorectal cancer, CRC; renal cell carcinoma, RCC; liver cancer, LIHC; lung cancer, LC; oral squamous cell carcinoma, OSCC; pancreatic cancer, PAAD; gastric cancer, GC; thyroid cancer, THCA.

We next asked whether this profound divergence would reshape the resulting peptide landscape. Among peptides with high HLA binding affinity (half maximal inhibitory concentration, IC_50_ ≤ 500 nM; **Methods**), only 18.2% of predicted microbial HLA-binding peptides were shared between tumor and peritumor tissues (**Fig. 1E**), ranging from 0.8% in RCC to 16.5% in lung cancer (LC) (**Extended Data Fig. S6**). Even when restricting to the 2,455 species resident in both tissue types, merely 18.6% of predicted HLA-binding peptides were shared (**Fig. 1F**), with proportions ranging from 0.2% in CESC to 16.8% in LC (**Extended Data Fig. S7**). The robustness of these findings was supported by saturation analysis, which confirmed that microbial species detection from our WTS data had reached plateaus (**Methods**; **Fig. 1G** and **Extended Data Fig. S8**), and mass spectrometry-based immunopeptidomics confirmed the physical presentation of predicted microbial peptides on MHC molecules (**Methods**; **Extended Data Table S4**).

Pairwise cosine similarity analysis showed that tumor–tumor comparison pairs exhibited significantly higher similarity in predicted HLA-binding peptide repertoires than peritumor-peritumor comparisons (**Methods**; *p* = 2.84 × 10⁻^5^, one-tailed Mann–Whitney test; **Fig. 1H**). Among tumor pairs, the highest similarity was observed between liver cancer (LIHC) and BLCA (cosine similarity = 0.42), while the highest peritumor– peritumor similarity occurred between LIHC and gastric cancer (cosine similarity = 0.19). Cross-cancer analysis further identified more than 15,000 shared predicted microbial HLA-binding peptides among liver, gastric, bladder, lung, and breast cancers (**Fig. 1I**).

### The Microbial Transcriptional Reprogramming Index (MiTRI) quantifies adaptive niche specialization

The profound transcriptional divergence between tumor and peritumor tissues suggests that resident microbes undergo MTR in response to the tumor microenvironment. Microorganisms, particularly bacteria, adapt to distinct environmental niches through selective gene expression ^23,24^. To quantify this adaptation in the tumor environment (TME), we compared the global transcriptome-wide distribution patterns of identical bacteria present in both tumor and peritumor samples (**Methods**). RNA sequencing (RNA-seq) coverage analysis of literature-reported tumor-associated bacteria revealed pronounced divergence among tumor and peritumor tissues ^25–30^ (**Extended Data Table S5** and **Fig. 2A–C**). *Fusobacterium nucleatum* showed markedly higher coverage in CRC tumor tissues than in matched peritumor tissues (*p* = 7.15 × 10⁻¹², one-tailed Mann– Whitney test; **Fig. 2A**). Similarly, *Cutibacterium acnes* was significantly enriched in breast cancer (BRCA) tumor tissues compared to peritumor tissue samples (*p* = 2.84 × 10⁻²⁰; **Fig. 2B**). Conversely, the probiotic *Bifidobacterium longum* displayed significantly reduced transcriptional coverage in BRCA tumor tissues relative to peritumor samples (*p* = 2.10 × 10⁻²⁷; **Fig. 2C**).

**Fig. 2.**
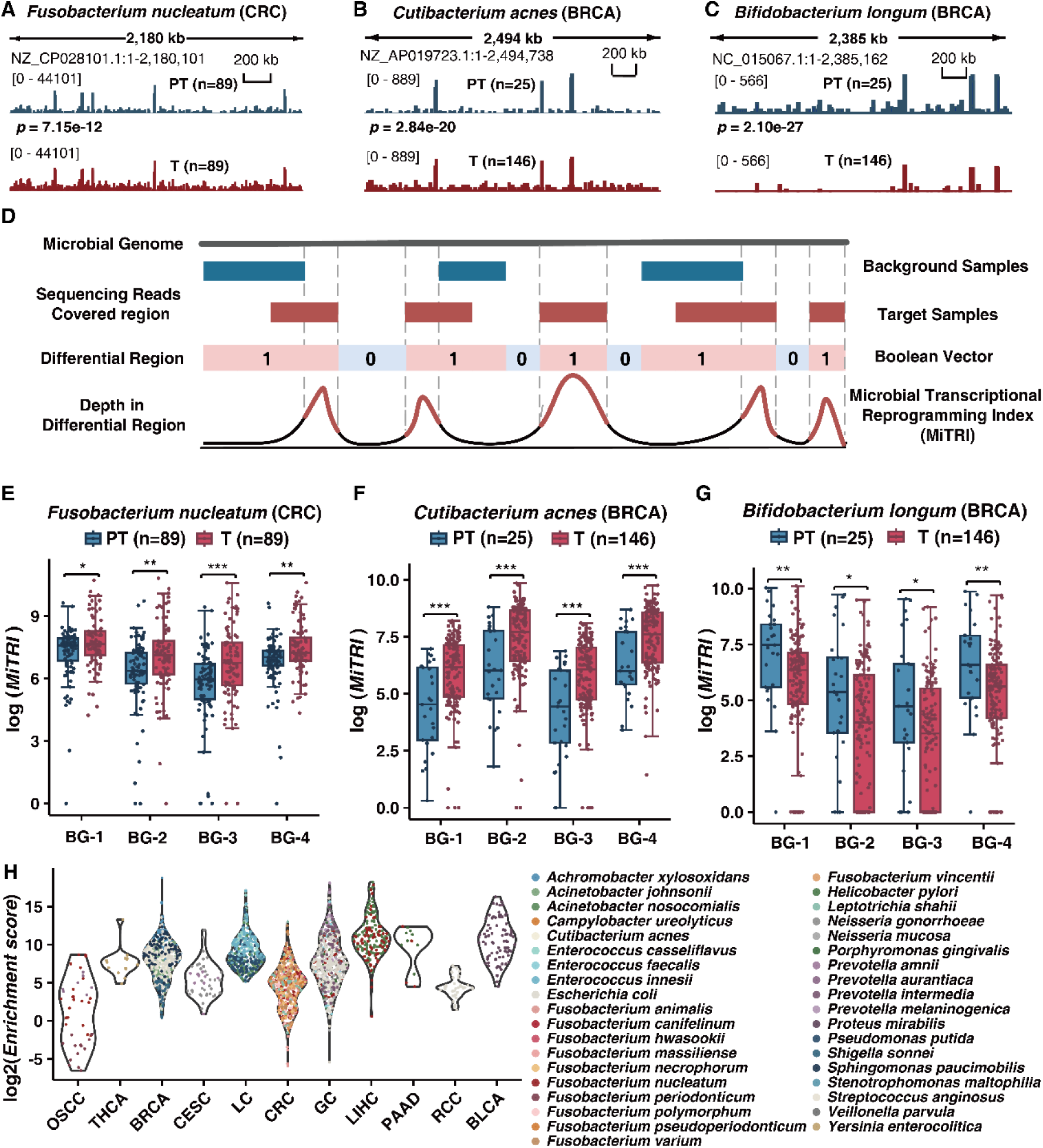
Distinct MTR patterns in tissue-associated microbes. **(A)** Genome-wide transcriptional RNA-seq coverage of *Fusobacterium nucleatum* in CRC tumor tissues (T; red; n = 89) and matched peritumor tissue samples (PT; blue; n = 89), showing significantly higher coverage in tumor tissues (*p* = 7.15 × 10⁻¹², one-tailed Mann– Whitney test). **(B)** Genome-wide transcriptional RNA-seq coverage of *Cutibacterium acnes* in BRCA tumor tissues (T; red; n = 146) and peritumor tissues (PT; blue; n = 25), with significantly higher coverage in tumor tissue samples (*p* = 2.84 × 10⁻²⁰, one-tailed Mann–Whitney test). **(C)** Genome-wide transcriptional RNA-seq coverage of *Bifidobacterium longum* in BRCA tumor tissues (T; red; n = 146) and peritumor tissues (PT; blue; n = 146), with significantly higher coverage in peritumor tissue samples (*p* = 2.10 × 10⁻²⁷, one-tailed Mann–Whitney test). **(D)** Workflow of calculating the Microbial Transcriptional Reprogramming Index (MiTRI). RNA-seq coverage of target (red) and background (blue) samples is mapped to the microbial genome. A Boolean vector records differential programming regions, where 1 denotes regions covered in target or background samples, and 0 denotes regions with no coverage in either. MiTRI is calculated by summing the coverage within the target-specific MTR regions (only covered in target samples), normalized by microbial read proportion and the length of the microbial transcriptional programming regions (labeled with 1). Higher MiTRI indicate stronger transcriptional adaptation to the target environment. **(E)** MiTRI values of *Fusobacterium nucleatum* in CRC across four background sets. The bacterium shows significantly stronger adaptation in tumor tissues (red; n = 89) compared to matched peritumor tissue samples (blue; n = 89). MiTRI values were log_e_-transformed for visualization. (*** *p* < 0.001, ** *p* < 0.01, * *p* < 0.05, Mann–Whitney test.) **(F)** MiTRI values of *Cutibacterium acnes* in BRCA across four background sets. The bacterium shows significantly stronger adaptation in tumor tissues (red; n = 146) compared to peritumor tissue samples (blue; n = 25). **(G)** MiTRI values of *B. longum* in BRCA across four background sets. The bacterium shows significantly stronger adaptation in peritumor tissues (blue; n = 25) compared to tumor tissues (red; n = 146). **(H)** Distribution of transcriptional enrichment scores (log2-transformed) for tumor-associated bacteria across 11 cancer types. Each point represents one sample-bacteria pair with statistically significant differential transcription (two-sided binomial test, p < 0.05). (Enrichment scores were calculated as the ratio of observed-to-expected read density in MTR regions, where expected density assumes uniform genome-wide distribution; Enrichment score > 1 indicate selective enrichment of MTR regions beyond genome-wide background.) Abbreviations: background, BG; 17 healthy brain samples, BG-1; nine glioblastoma peritumor tissue samples, BG-2; 17 healthy brain and nine glioblastoma peritumor tissue samples, BG-3; and six healthy testis samples, BG-4.

Having established that changes driven by environmental pressure are detectable by RNA-seq data, we developed the Microbial Transcriptomic Reprogramming Index (MiTRI) to formally measure the degree of MTR between target and background samples (**Methods; Fig. 2D**). We defined the MTR regions as genomic loci that were exclusively transcribed in target samples but not in background samples, identified at single-nucleotide resolution. For comparing the MiTRI between tumor and peritumor tissues, we selected tissues with naturally low microbial loads, such as brain and testis ^31–36^, as the background to establish baseline transcriptional states. Application of MiTRI across tumor-associated bacteria confirmed transcriptional reprogramming patterns. *Fusobacterium nucleatum* and *Cutibacterium acnes* displayed significantly higher MiTRI in tumor tissues compared to peritumor tissues across all background choices (**Fig. 2E–F**), whereas *Bifidobacterium longum* exhibited the inverse pattern (**Fig. 2G**). Additional tumor-associated bacteria similarly showed higher MiTRI in tumor tissues relative to peritumor tissues, such as *Helicobacter pylori*, *Porphyromonas gingivalis*, *Prevotella intermedia*, and *Enterococcus faecalis* (**Extended Data Table S5** and **Extended Data Fig. S9–S12**).

To demonstrate tumor specificity, we explored tumor-specific MTR regions for each bacterium in each tumor sample using peritumor tissues of the corresponding cancer type as background (**Methods**). We calculated transcriptional enrichment within these regions to evaluate whether they were microbial selectively activated loci (**Methods**). Among sample-bacteria pairs showing significant differential transcription (binomial test, *p* < 0.05; **Methods**; **Extended Data Table S5**), 97.4% of sample-bacteria pairs exhibited transcriptional enrichment within tumor-specific MTR regions (enrichment score > 1; **Methods**; **Fig. 2H**) .

### Tumor-specific MTR expands the immunogenic peptide repertoires in tumor tissues

Having established that resident microbes undergo MTR in the TME, we investigated whether this MTR process generates tumor-specific immunogenic peptides that could serve as safe therapeutic targets. We focused on the tumor-associated bacteria characterized above and examined wheather tumor-specific MiTRI relates to peptide generation and immunogenicity (**Extended Data Table S5; Methods**).

We observed tumor-specific MiTRI positively correlated with microbial abundance in tumor tissues( Spearman’s *ρ* = 0.46, *p* < 2.2 × 10^−16^) and with the number of distinct microbial genes specificly expressed in tumor tissues (Spearman’s *ρ* = 0.26, *p* < 2.2 × 10⁻¹⁶; **Fig. 3A; Methods**). Critically, tumor-specific MiTRI significantly correlated with the number of tumor-specific predicted HLA-binding peptides (Spearman’s *ρ* = 0.22, *p* = 1.8 × 10^−1^^4^; **Fig. 3A; Methods**). We next evaluated the immunogenic quality of peptides derived from these tumor-specific MTR regions. These peptides demonstrated superior HLA-binding affinity (mean IC_50_ = 12.97 nM) compared with peptides from entire bacterial genomes (mean IC_50_ = 19.25 nM; **Extended Data Fig. S13**). Moreover, tumor-specific MiTRI positively correlated with the predicted immunogenicity of peptides from tumor-specific MTR regions (Spearman’s *ρ* = 0.29, *p* < 2.2 × 10⁻¹⁶; **Fig. 3A**). The positive correlation between tumor-specific MiTRI and immunogenicity was replicated using two additional algorithms: BigMHC (Spearman’s *ρ* = 0.13, *p* < 3.4 × 10^−6^; **Extended Data Fig. S14**) and DeepImmuno (Spearman’s *ρ* = 0.21, *p* = 8.0 × 10^−1^^3^; **Extended Data Fig. _S14_**_) 37,38._

**Fig. 3.**
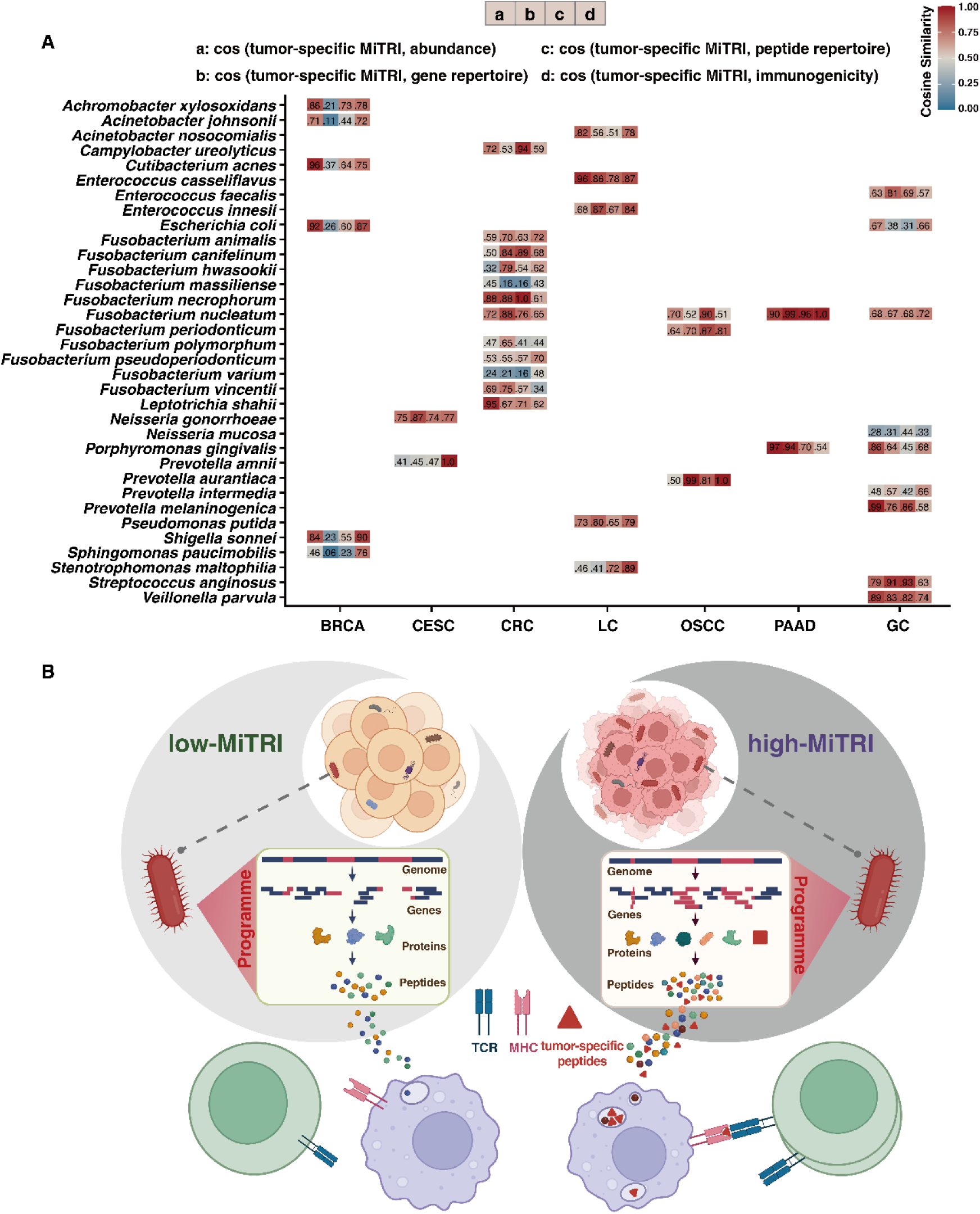
Correlation of MTR with enhanced microbiome diversity in tumors. **(A)** Correlations between tumor-specific MiTRI and microbial abundance in tumor tissues (a), number of distinct genes exclusively expressed in tumor tissues(gene repertoires, b), number of distinct tumor-specific predicted HLA-binding peptides (peptide repertoire, c), and immunogenicity of peptides from tumor-specific MTR regions (d).(cosine similarity: 0.28–0.99, 0.06–0.99, 0.16–1.00, and 0.33–1.00, respectively). **(B)** Schematic illustration of the proposed hypothesis: MTR drives the generation of tumor-specific peptides, expanding the reservoir of more immunogenic, tumor-specific microbial antigens.

To validate whether predicted microbial peptides are physically presented by MHC molecules, we performed mass spectrometry-based immunopeptidomics and detected 71 microbial peptides derived from 31 species on an CRC recurrent metastasis specimen (70 presented by MHC class I, 1 by MHC class II; **Methods**; **Extended Data Table S4**). The MHC class II-presented peptide detection is consistent with tumor cell MHC class II expression demonstrated in CRC organoids, patient-derived immunopeptidomics, and mechanistic validation of spatial transcriptomics ^2,3,39–42^. Among 20 HLA-binding peptides (19 MHC-I-presented, 1 MHC-II-presented; IC_50_ ≤ 500 nM), 5 (25%, 5/20; 4 MHC-I-presented, 1 MHC-II-presented) derived from tumor-specific MTR regions (**Extended Data Table S4** and **Extended Data Fig. S15**). Collectively, these pan-cancer analyses and immunopeptidomics establish MTR as a principle that enlarges the detectable pool of immunogenic, tumor-specific microbial peptides presented by MHC (**Fig. 3B**), enhancing the tumor peptide landscape and immune recognition within the TME.

### MTR Correlated with treatment response in breast and colorectal cancers

To establish the clinical significance of MTR, we investigated its association with treatment response in BRCA and CRC, two types of cancers with well-characterized microbial enrichment^14,43,44^. In pre-treatment BRCA tissues, patients who responded to neoadjuvant therapy exhibited significantly higher MiTRI values for the tumor-associated bacteria *Cutibacterium acnes* than non-responders (*p* < 0.01 to *p* = 0.03, **Extended Data Fig. S16**).

To further investigate this relationship, we performed multi-omics profiling on MSS-CRC patients treated with neoadjuvant therapy, a setting where conventional neoantigens are scarce and microbial contributions are likely pronounced (**Methods**; **Fig. 4A–B**). Whole-exome sequencing data confirmed genomic homogeneity across samples, with driver mutations showing high concordance between responders and non-responders (Jaccard indices: 0.88–1.00, **Fig. 4C** and **Extended Data Table S6**) ^45^, consistent with the limited neoantigen load characteristic of MSS-CRC ^7^. Strikingly, however, principal component analysis of host transcriptomes revealed a clear separation between the two groups (**Fig. 4D**).

**Fig. 4.**
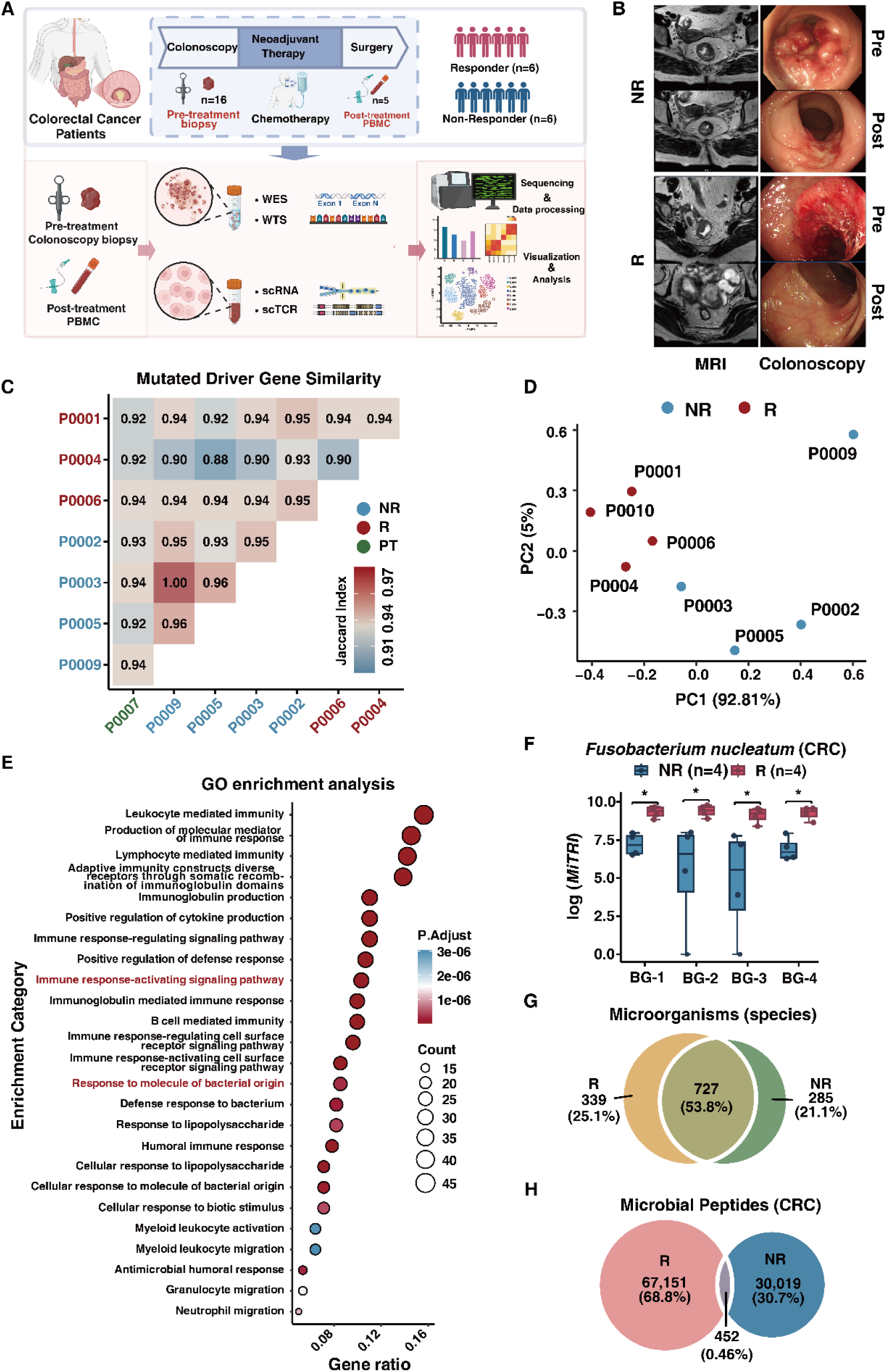
MTR and its association with chemotherapy response in CRC patients. **(A)** Overview of the study design comparing CRC samples from chemotherapy responder (R) and non-responder (NR) group. **(B)** Representative MRI and colonoscopy images of pre- and post-treatment for R and NR patients. **(C)** Jaccard similarity of mutated driver genes across patient groups (R, red, n = 3; NR, blue, n = 4; peritumor (PT), green, n = 1). **(D)** Principal component analysis (PCA) of differential gene expression in pre-treatment CRC samples, showing different clustering of R (red; n = 4) and NR (blue; n = 4) patients. **(E)** Gene Ontology (GO) enrichment analysis of upregulated genes in the responder group. Significantly enriched GO terms include “immune response-activating signaling pathway” and “response to molecule of bacterial origin.” **(F)** Comparison of *Fusobacterium nucleatum* MiTRI between R and NR groups (* *p* < 0.05, Mann–Whitney test). **(G)** Venn diagram showing the overlap of microbial species between R and NR groups. **(H)** Venn diagram showing the overlap of predicted microbial HLA-binding peptide repertoires between R and NR groups.

This transcriptional divergence was functionally anchored in the host immune response to microbes. GO enrichment analysis of upregulated host genes highlighted significant enrichment of immune pathways in the responders, including the “immune response-activating signaling pathway” and “response to molecule of bacterial origin” (**Fig. 4E**). Corroborating this, both lipopolysaccharide staining of gram-negative bacteria and WTS-based quantification of CRC-associated bacteria confirmed differential microbial abundance in responder groups (**Extended Data Fig. S17–S18**). Crucially, these bacteria exhibited significantly higher MiTRI values in the responder group (**Fig. 4F** and **Extended Data Fig. S19**), corroborated by their transcriptional read distribution patterns (**Extended Data Fig. S20**). Notably, despite the overall microbial compositions showed 53.8% species resident in responder and non-responder groups (**Fig. 4G**), their number of predicted HLA-binding peptides showed minimal sharing at only 0.46% (**Fig. 4H**).

### MTR-derived tumor-specific epitopes drive a dominate tumor-reactive T cell response

We next asked whether MTR-derived microbial peptides contribute directly to anti-tumor immunity. Profiling of pre-treatment tumor tissues showed comparable baseline immune infiltration between future responders and non-responders (**Methods**; **Fig. 5A**). Following neoadjuvant therapy, responders achieved complete pathological response, precluding post-treatment tumor analysis (**Fig. 4B**). We therefore turned to peripheral blood mononuclear cells (PBMCs) and performed single-cell RNA sequencing (scRNA-seq) to track systemic immune dynamics ^46^. In responders, we observed profound immune remodeling after treatment, characterized by a marked expansion of effector T cell populations (CD8⁺ Temra/Teff, and Tem) and γδ T cells, accompanied by decreased CD4⁺ Temra/Teff, CD4⁺ naive/central memory T cells (Tn/Tcm) and CD4⁺ Treg cells (**Methods**; **Fig. 5B** and **Extended Data Fig. S21**; **Extended Data Table S7**), consistent with therapy-induced immune activation.

**Fig. 5.**
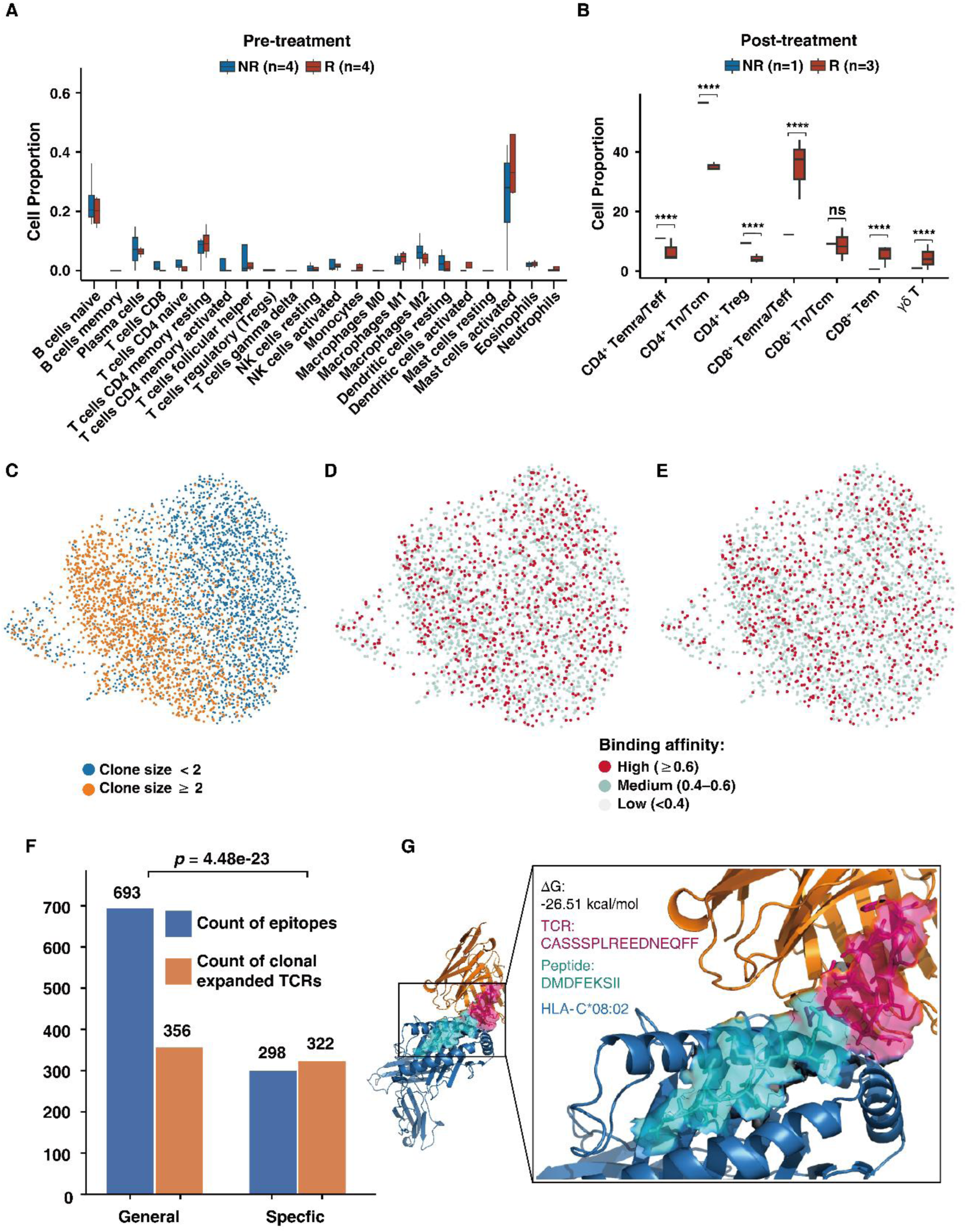
MTR-derived epitopes show enhanced recognition by TCRs with large clone sizes in therapy responders. **(A)** Immune cell composition in pre-treatment tumor samples showed no significant differences between chemotherapy responder (R, n = 4) and non-responder (NR, n = 4) groups, as inferred from deconvolution of bulk RNA-seq data using CIBERSORT with LM22 signature matrix (no significant difference, Mann– Whitney test). **(B)** scRNA analysis of post-treatment PBMCs reveals distinct immune profiles between R (n = 3) and NR (n = 1) groups. (**** *p* < 0.0001, ** *p* < 0.01, ns *p* ≥ 0.05, chi-square test). **(C)** UMAP visualization of TCR clonotypes from single-cell TCR sequencing data, colored by clone size: expanded clones (clone size ≥2, orange) and non-expanded clones (clone size <2, blue). **(D)** UMAP visualization of expanded TCRs with high predicted TCR-binding affinity (≥ 0.6) recognizing general candidate epitopes derived from whole-genome regions of nine CRC-associated bacteria. **(E)** UMAP visualization of expanded TCRs with high predicted TCR-binding affinity (≥ 0.6) recognizing specific candidate epitopes derived from tumor-specific MTR regions of nine CRC-associated bacteria. **(F)** Comparison of TCR recognition frequency between general epitopes (whole-genome derived) and specific epitopes (MTR region-derived) from nine CRC-associated bacteria. Specific epitopes (n=298) were recognized by 322 TCRs (1.08 TCRs per epitope), while general epitopes (n=693) were recognized by 356 TCRs (0.51 TCRs per epitope), demonstrating a 2.10-fold higher recognition rate for MTR-derived epitopes (p = 4.48 × 10⁻²³, Poisson rate ratio test).**(G)** Structural model of the HLA–epitope–TCR complex showing the β-chain CDR3 loop (CASSSPLREEDNEQFF) engaging the microbial candidate epitope DMDFEKSII presented by HLA-C*08:02. The interaction interface is characterized by van der Waals contacts, hydrogen bonds, and salt bridges, primarily mediated by the TCR β-chain. Surface rendering illustrates a contiguous binding surface with shape and charge complementarity, consistent with the negative binding free energy (ΔG = −26.51 kcal/mol).

To directly test whether this activated T cell response targeted MTR-derived epitopes, we integrated pre-treatment microbial transcriptomes with post-treatment single-cell TCR sequencing (scTCR-seq). From a representative responder, we identified 1,504 predicted HLA-binding peptides derived from 37 bacteria in a responder sample (IC_50_ ≤ 500 nM;**Extended Data Table S8**). TCR-binding affinity predictions indicated that that 78.1% (1,174/1,504) of these peptides were likely recognized by TCRs with high binding affinity (≥ 0.6). Among these, 45.6% (529/1,174) originated from nine known CRC-associated bacteria that were transcriptionally reprogrammed (higher MiTRI) in responders, and 41.6% (220/529) of these were encoded exclusively within tumor-specific MTR regions..

We then quantified the immunodominance of MTR-derived epitopes by analyzing T cell clonal expansion (**Fig. 5C**). Among 597 high-affinity TCR clonotypes, 53 underwent post-treatment expansion (clone size ≥ 2). The vast majority of these expanded clonotypes (42/53) recognized epitopes from the nine MiTRI-high, CRC-associated bacteria(**Fig. 5D**), and 79.1% (34/43) were specific for epitopes derived from tumor-specific MTR regions (**Fig. 5E** and **Extended Data Table S8**). This bias toward MTR regions was striking. Although MTR-derived epitopes constituted a smaller fraction of the total epitope pool, they attracted a disproportionate share of T cell attention. The TCR recognition rate for MTR-derived epitopes was 2.10-fold higher than that for epitopes derived from the global microbial genome (*p* = 4.48 × 10⁻^23^, Poisson rate ratio test; **Fig. 5F**).

To provide a structural basis for this enhanced immunogenicity, we modeled the HLA-epitope-TCR complexes. Those involving MTR-derived epitopes and their cognate, expanded TCRs exhibited significantly lower binding free energies (ΔG)(**Extended Data Fig. S22** and **Extended Data Table S9**). These biophysical data solidify the conclusion that MTR generates epitopes which are structurally optimized for effective T cell engagement (**Fig. 5G** and **Extended Data Fig. S23**).

## Discussion

Previous investigations of tumor-associated microbiomes have primarily focused on cataloging compositionally enriched species and their roles in tumor progression^13–15^. While these studies successfully identified which microbes colonize tumors and demonstrated that their peptides can activate anti-tumor immunity, a general principle to ensure the safe targeting of microbial antigen has remained elusive. Our pan-cancer study resolved this safety gap by demonstrating that compositionally similar microbial communities in tumor and peritumor tissues give rise to distinct peptide repertoires through MTR, thereby providing a theoretical foundation for the safe clinical deployment of microbiome-directed immunotherapy.

We discovered that identical microbial species present in both tumor and adjacent normal tissues can generate tumor-specific peptides. This specificity originates from genes that are selectively activated within the TME ^47^, a process driven by tumor-induced transcriptional reprogramming.. Supporting this, GO enrichment analysis of microbial transcriptomes across 2,455 species resident in both tissue types revealed starkly divergent functional states: 491 terms significantly enriched in tumor tissues compared to 244 in peritumor tissues, with only five overlapping terms (FDR-BH < 0.05). This demonstrates that identical microbial species exhibit fundamentally different transcriptional programs in direct response to their tissue niche.

The tumor-specific transcriptional activation of microbial genes provides the essential protein reservoir for generating tumor-associated microbial peptides. This notion is supported by our observation that microbial peptides derived from tumor tissues are more prevalent than those from peritumor tissues. Furthermore, proteins encoded within MTR regions, owing to their selective expression under tumor conditions, are prime candidates for enhanced immunogenicity. Consistently, our findings demonstrate that MTR-derived peptides display superior predicted HLA-binding affinity compared to those from non-MTR regions. Together, these results establish a mechanistic and theoretical basis for identifying tumor-specific microbial peptides guided by the MTR principle.

To operationalize this principle, we developed MiTRI as a quantitative metric of microbial adaptation to the tumor niche. A higher MiTRI value indicates greater degree of transcriptional reprogramming under tumor-specific pressures, a pattern strongly exhibited by known tumor-associated microbes. Crucially, microbes with elevated MiTRI values produced more tumor-specific peptides with enhanced predicted immunogenicity. These findings demonstrate the MiTRI framework as a robust and feasible tool for prioritizing therapeutic targets,providing a methodological foundation for discovering broad-spectrum tumor-specific peptides.

Beyond individual cancer types, the identification of over 15,000 shared microbial peptides across five major tumor types suggests the existence of conserved reprogramming patterns, opening the door to pan-cancer therapeutic strategies. Unlike patient-specific tumor neoantigens requiring personalized approaches, MTR-derived epitopes occupy a therapeutic sweet spot: they are tumor-specific yet shared across patients. This could enable off-the-shelf vaccines or adoptive cell therapies targeting conserved microbial reprogramming signatures. The consistency of MTR across diverse malignancies suggests convergent evolution of microbial adaptation strategies, potentially reflecting fundamental constraints on how microbes survive in hostile tumor microenvironments.

Our findings in MSS-CRC provide compelling clinical proof-of-concept for this principle in a therapeutically challenging context. MSS-CRC exhibits low tumor neoantigen burden, creating an urgent need for alternative immunotherapy targets. We observed that neo-adjuvent therapy responders exhibited significantly higher MiTRI for reported CRC-associated bacteria compared to non-responders, coupled with the 2.10-fold enhanced TCR recognition of MTR-derived epitopes compared to genome-wide epitopes. Remarkably, 79.1% of clonally expanded TCRs were predicted to recognize MTR-derived epitopes, indicating these epitopes are associated with therapeutically meaningful responses rather than representing bystander activation. These human data extend recent mouse work demonstrating that killing bacteria can release microbial therapeutic targets, providing clinical evidence that MTR actively generates immunotherapy-relevant peptides in cancer patients^6^.

In conclusion, our work redefines the understanding of tumor-microbe dynamics by quantifying that the transcriptional state of the microbiome, rather than its mere composition, dictates tumor-specific antigenicity. By systematically characterizing MTR across cancer types and linking it to immunogenic epitopes, we provide both a biological insight and an actionable principle for therapeutic development. As microbiome-based immunotherapies advance toward clinical implementation, incorporating transcriptional state analysis through MiTRI will be paramount for intelligent target prioritization and precise patient stratification. This principle finally enables the safe harnessing of the microbiome’s therapeutic potential, minimizing the risk of on-target, off-tumor toxicity against commensal microbiota.

## Methods

### Sample Collection and Data Acquisition

#### Patient Enrollment

This study enrolled patients with advanced-stage microsatellite-stable (MSS) colorectal cancer (CRC) defined as American Joint Committee on Cancer (AJCC) stage II-III disease (specifically T2 with nodal involvement [N+] or ≥ T3 lesions, all M0). Treatment involved administering 85 mg/m² of a platinum-based agent and 400 mg/m² of leucovorin intravenously on day 1, followed by a bolus of 400 mg/m² of a fluoropyrimidine-based drug and a continuous infusion of 2400 mg/m² of the fluoropyrimidine-based drug over 46 to 48 hours. This regimen was repeated administered every 2 to 3 weeks for a total of 4 to 6 cycles. Patients were classified as responsive to neoadjuvant therapy if at least two of the following criteria were met: (1) No palpable tumor mass in the original tumor site upon digital rectal examination; (2) No visible tumor under endoscopy, or only minor superficial ulcers/scars; (3) Evidence of complete response (CR) or extensive fibrotic changes with minimal residual lesions, classified as MRI diagnostic grade 1-2; (4) Pathological tumor regression grade (TRG) of 0–1, or partially 2 (with a fibrotic component close to 50%).

#### Specimen Collection

Pre-treatment biopsy samples were collected using endoscopic forceps to obtain tumor tissue. Depending on tumor size, 2 to 3 pieces (approximately 2 mm in diameter) were excised. Specimens were rinsed with saline to remove blood and mucus, immediately snap-frozen on dry ice for 2 hours during transportation, and stored at −80°C. All procedures were approved by the Ethics Committee of the Hefei Cancer Hospital, Chinese Academy of Sciences. Written informed consent was obtained from all participants before sample collection.

#### Public Sequencing Data Collection

Transcriptome sequencing data were collected from NCBI Sequence Read Archive (SRA) database. The focus was on solid tumors, including bladder, breast, cervical, colorectal, kidney, liver, lung, oral, pancreas, gastric, and thyroid cancers. Data collection criteria included the use of whole-transcriptome sequencing with random priming following ribosomal RNA depletion. Datasets for each cancer type needed to include peritumor tissue from healthy individuals or peritumor tissue adjacent to tumors as controls. After excluding samples with a low microbial reads proportion (<10^−6^), a total of 1,868 samples spanning 11 cancer types were collected (Extended Data Table S2). The dataset comprised tumor tissues and peritumor tissues, with some projects providing patient-matched pairs while others included unmatched samples from the same cancer type.

### Sequencing for In-house Samples

#### Whole Exome Sequencing

DNA extraction was performed using the HiPure Universal DNA Kit (Catalog Number: D3018-02, Magen). For library preparation, the Hieff NGS^®^ OnePot Pro DNA Library Prep Kit for Illumina**^®^** (Catalog Number: 12205, Yeasen) was used. Sequencing was conducted on the DNBSEQ-T7 platform, producing 150 bp paired-end reads.

#### Whole Transcriptome Sequencing

Whole transcriptome sequencing (WTS) was carried out using a strand-specific strategy on the BGI platform. Total RNA was extracted using the Hieff NGS ^®^ MaxUp Human rRNA Depletion Kit (rRNA & ITS/ETS) (Catalog Number:12257ES24, Yeasen). RNA integrity was assessed using the Agilent 2100 Bioanalyzer (Agilent Technologies). Library preparation was conducted using the Hieff NGS^®^Ultima Dual-mode mRNA Library Prep Kit (Catalog Number: 12301ES96, Yeasen). Sequencing was performed on the DNBSEQ-T7 platform, generating 150 bp paired-end reads.

#### Single-cell RNA and Single-cell TCR Sequencing

Peripheral blood mononuclear cells (PBMCs) were isolated from the blood of responders and non-responders to neoadjuvant therapy using density centrifugation over Ficoll-Paque plus (Catalog Number: 04-03-9391/03, stemcell) and washed 3 times with PBS. Single-cell RNA libraries were prepared using the Chromium Single Cell 5’v2 Reagent (10x Genomics), and Chromium Single Cell V(D)J Reagent kits (10x Genomics) for single-cell RNA and TCR libraries. Each sequencing library was generated with a unique sample index. Sequencing libraries were quantified using a High Sensitivity DNA Chip (Agilent) on a Bioanalyzer 2100 and the Qubit High Sensitivity DNA Assay (Thermo Fisher Scientific) and sequenced on DNBSEQ-T7 (MGI Tech Co., Ltd.) with PE150 read length.

#### Immunohistochemistry

The paraffin-embedded tissue sections fixed with formalin were cut into 5 μM slides. The slides were baked at 60°C for 2 hours, then dewaxed with xylene and rehydrated with ethanol for 3 minutes (100% −90% −80%). Antigen retrieval was performed in sodium citrate antigen retrieval solution (Solarbio, # C1032) using a microwave oven, maintaining sub-boiling temperature for 15 minutes, and then allowed to cool to room temperature naturally. The slices were blocked with 3% hydrogen peroxide (SCR, #10011218) at room temperature for 15 minutes, followed by blocking with 10% BSA (Solarbio, #A8020) at 37°C for 30 minutes. The primary antibody (anti-LPS, Hycult Biotech, #HM6011-20UG, 1:200) was added to the slides and incubated overnight at 4°C. The next day, the slides were washed with PBS and then incubated with the secondary antibody (goat anti-mouse IgG secondary antibody, Abkine, #A21010, 1:500) at room temperature for 1 hour. DAB staining was performed using the DAB staining kit (ZSGB-Bio, #ZLI-9018), followed by hematoxylin counterstaining. The slides were dehydrated with ethanol for 3 minutes (80% −90% −100%) and mounted with neutral gum. Immunohistochemical images were captured using a microscope (YUESHI, China) at 10X and 20X magnifications.

### Mass spectrometry-based immunopeptidomics validation

#### MHC-I and MHC-II peptide Enrichment

MHC-I and MHC-II peptides were obtained from human tissue as described previously^48,49^. In each group, around 300 mg CRC tissue were used for immunopeptides isolation. CRC tissues were homogenized in lysis buffer with 0.25% sodium deoxycholate, 1% n-octyl glucoside, 100 mM PMSF, 0.2 mM iodoacetamide, and protease inhibitor cocktail in Gibco’s Dulbecco’s phosphate-buffered saline (DPBS). Lysate was further cleared by centrifugation for 50 min at 17,000g at 4 °C. The MHC-I was purified with W6/32 antibody covalently bound to Protein-A Sepharose beads from supernatant, while MHC-II was purified with IVA12 antibody covalently bound to Protein-G Sepharose beads. Beads were then washed with buffer A (150 mM NaCl, 20 mM Tris HCl) and 400 mM NaCl, 20 mM Tris HCl. The MHC-I and MHC-II complexes were eluted with 10% acetic acid. Eluate was then loaded on Sep-Pak tC18 cartridges (Waters, 100 mg) and washed with 0.1% TFA and 2% ACN in 0.1% TFA, sequentially. The MHC-I peptides were separated from MHC-I complexes on the tC18 cartridges by eluting with 28% ACN in 0.1% TFA. And the MHC-II peptides were eluted from tC18 cartridges by 32% ACN in 0.1% TFA. The purified immunopeptides were dried using vacuum centrifugation.

#### Immunopeptides Sequencing with data-independent acquisition

The samples were sequenced using an Orbitrap Astral mass spectrometer (Thermo Fisher Scientific) coupled to a Vanquish HPLC system (Thermo Fisher Scientific). Briefly, the dried immunopeptides were resuspended in 12 μL of 2% ACN in 0.1% TFA (Trifluoroacetic acid). Two replicates of 5 μL samples injected to mass spectrometer. Each sample was separated on an Aurora Elite C18 column (15 cm x 75 um lD, 1.7 um) at a flow rate of 450 nL/min by 25 min gradient of buffer A (0.1% formic acid) and buffer B (80% ACN and 0.1% formic acid). The peptides were ionized by 1.9K spray voltage followed by data-independent acquisition (DIA) analysis. A full MS1 scan was acquired from 400 to 900 m/z with a resolution of 240,000 and the ion accumulation time of 20 ms, followed by 40 DIA MS2 scans acquired from 150 and 2000 m/z with 10 m/z isolation windows and an automatic gain control (AGC) of 5×10^6^. A stepped normalized collision energy (30) was employed, and the maximum ion accumulation was set to auto. FAIMS voltage was turned on and CV value was set to −48.

The DIANN software package (version 1.8) was used for analyzing MS files^50^. To reveal the microbe-derived immunopeptides, we generated the spectral library against microbe immunopeptide database and the UniProt database (human-reviewed sequences Release 2025_03). The raw DIA MS files were directly loaded in the DIANN workflow and searched against. The peptide length range was set to 8-15 for MHC-Ⅰ MS data search and 12-20 for MHC-Ⅱ MS data search. The other parameters were set as default. For peptide quantification and filtering, peptide intensities from two technical replicates were averaged. The averaged intensities were then log2-transformed (adding a pseudocount of 1 to avoid undefined values). Only peptides with 𝑙𝑜𝑔2(𝑖𝑛𝑡𝑒𝑛𝑠𝑖𝑡𝑦 + 1)) > 10 were retained for downstream analysis. The predicted spectra for representative immunopeptides were generated using Prosit for comparison with experimental DIA spectra^51^.

### Construction of Human-Associated Microbial Reference Dataset

#### Refinement of Human-Associated Microbial Taxa

A curated and updated reference dataset of tumor-associated microbes was constructed, based on a published dataset^13^, and further refined with data from the Human Microbiome Project (HMP) ^21^ and additional literature sources (Extended Data Fig. S1, Extended Data Table S1). This resulted in a comprehensive microbial list consisting of 355 bacterial genus. The taxonomic hierarchy was determined using NCBI taxonomy IDs sourced from the NCBI taxonomy tree (ftp://ftp.ncbi.nlm.nih.gov/pub/taxonomy/taxdump.tar.gz, updated Nov. 7, 2024). In total, 8,315 high-quality genome sequences at the strain or species level were selected for inclusion in the dataset, using assembly summary data (ftp://ftp.ncbi.nlm.nih.gov/genomes/refseq/assembly_summary_refseq.txt, updated Nov. 14, 2024).

#### Manual Curation of Tumor-Associated Microbial Taxa

After the initial refinement, a manual review was conducted to refine the list, focusing on taxa that were most relevant to tumorigenesis. The microbial list was further curated to ensure its applicability to cancer research, with a final selection of 3,199 high-quality bacterial taxa and 1,262 viral taxa (Extended Data Table S1). To enhance this curation, the large language model, ChatGPT 4.0^52^, was employed to assess each microbial taxon based on its potential to colonize human tissues, association with tumorigenesis, and its impact on anti-tumor treatments. The prompt used for this task was as follows: “The list below contains microbiome taxa at the species or strain level. Please provide the following information for each: Brief Description, Potential to Colonize Human Tissues, Association with Tumorigenesis and Progression, and Impact on Anti-Tumor Treatments.” To ensure manageable output, taxa were randomized and submitted in batches of 50. Each GPT-generated response underwent manual validation by cross-referencing literature and expert evaluation.

#### Criteria for Microbial Classification

**Potential to Colonize Human Tissues**. Microbial taxa are classified based on their potential to establish and persist in human tissues, which is determined by factors such as microbial adherence mechanisms, invasiveness, and biofilm formation. Taxa are categorized into the following groups:

***Low:*** Taxa that rarely colonize human tissues or are found in minimal abundance.

***Moderate:*** Taxa that are occasionally found in specific human tissues but do not dominate the microbiome.

***High:*** Taxa that are frequently present and dominate specific tissues, indicating strong colonization potential.

***Require Manual Assessment:*** Taxa exhibiting indeterminate colonization patterns due to ambiguous metagenomic signatures or insufficient data resolution underwent manual verification. Classification was adjudicated through evidence synthesis from peer-reviewed literature and orthogonal experimental datasets.

**Association with Tumorigenesis and Progression**. Microbial taxa are assessed for their potential role in cancer development and progression. This includes evaluating evidence of inflammation, immune modulation, and carcinogenesis. Microbes are classified according to their established or suspected involvement in cancer processes:

***Tumorigenic*:** Taxa with strong evidence linking them to cancer development, often through chronic inflammation or other carcinogenic mechanisms.

***Opportunistic Pathogens:*** Taxa that do not directly cause cancer but may compromise the immune system or create a favorable environment for cancer progression.

***Probiotic or Beneficial Microbes:*** Taxa that may provide protective effects against cancer or enhance the effectiveness of cancer treatments.

***Require Manual Assessment:*** Taxa exhibiting indeterminate colonization patterns due to ambiguous metagenomic signatures or insufficient data resolution underwent manual verification. Classification was adjudicated through evidence synthesis from peer-reviewed literature and orthogonal experimental datasets.

**Impact on Anti-Tumor Treatments**. The impact of microbial taxa on cancer treatment outcomes is evaluated based on their influence on drug metabolism, immune responses, and overall treatment efficacy. The classification includes:

***Positive Impact:*** Taxa that enhance treatment efficacy, such as those producing compounds with anti-cancer activity or modulating the immune response to favor treatment.

***Negative Impact:*** Taxa that may interfere with treatment by altering drug metabolism or immune responses in a way that reduces the effectiveness of therapies.

***Neutral or Limited Impact:*** Taxa that have little or no observed effect on cancer treatments. ***Require Manual Assessment:*** Taxa exhibiting indeterminate colonization patterns due to ambiguous metagenomic signatures or insufficient data resolution underwent manual verification. Classification was adjudicated through evidence synthesis from peer-reviewed literature and orthogonal experimental datasets.

### Construction of Reference Genome and Protein Dataset

The final microbial genome sequences were downloaded from NCBI RefSeq (ftp://ftp.ncbi.nlm.nih.gov/refseq/release/bacteria/*.fna.gz and ftp://ftp.ncbi.nlm.nih.gov/refseq/release/viral/viral.1.1.genomic.fna.gz, updated Sep. 7, 2024). A microbial reference genome dataset was then constructed using BWA-MEM (v0.7.17) ^53^ and GATK *PathSeqBuildReferenceTaxonomy* (v4.6)^54^. To support downstream protein-level analyses, a custom protein dataset was generated based on the selected microbial genomes created with the *makeblastdb* tool from BLAST (v2.14.0)^55^, which obtained from NCBI (ftp://ftp.ncbi.nlm.nih.gov/ *genomes/all/*, updated on Sep. 7, 2024).

### Data Processing

#### Quantitative Host Gene Expression

RNA sequencing reads were aligned to the human reference genome (GRCh38) using STAR (v2.5.2b) with the following parameters^56^:

*--outSAMunmapped Within* (retain unmapped reads in the output),

-*-outFilterType BySJout* (enable junction-aware filtering),

*--outFilterMultimapNmax 20* (allow up to 20 mapping locations per read),

*--outFilterMismatchNmax 999* (permit up to 999 mismatches per read pair),

*--outFilterMismatchNoverLmax 0.04* (set mismatch density to ≤ 4% of read length),

*--alignIntronMin 20* and *--alignIntronMax 1,000,000* (define intron size constraints),

*--alignSJoverhangMin 8* (require a minimum of 8 nucleotides overhanging splice junctions),

*--quantMode TranscriptomeSAM* (output alignments in transcriptome coordinates). Gene expression quantification was performed using RSEM (v1.2.28) via the *rsem-calculate-expression*command with the following options:

*--paired-end* (paired-end reads),

*--alignments* (use precomputed STAR alignments),

*--p* (multithreaded computation),

*--no-bam-output* (suppress intermediate BAM generation).

### Identification and Annotation of Host Variants

Raw WES data were processed for quality control using Fastp (v0.23.2) to remove low-quality reads and adapter sequences^57^. Filtered reads were aligned to the human reference genome (GRCh38) with BWA-MEM ^58,59^ . The resulting SAM files were converted to sorted BAM format and indexed using SAMtools (v1.5)^60^. Duplicate reads were marked using GATK Spark MarkDuplicatesSpark. Base quality scores were recalibrated in two steps: generating recalibration models with BaseRecalibratorSpark using known variant resources from the Mills and 1000 Genomes Project^61^, dbSNP^62^, and the 1000 Genomes Phase 1 SNP dataset^63^, and then applying adjustments with ApplyBQSRSpark. Somatic variants were called using GATK Mutect2 on BQSR-processed BAM files, and candidate mutations were filtered to enhance reliability using FilterMutectCalls.

For variant annotation, we used VEP ^64^ within Singularity containers (v3.9.0-rc.3)^65^. The workflow utilized the GRCh38, with key parameters including offline mode with local cache, VCF output format, and transcript annotations (--symbol, --hgvs, --tsl, --biotype). Plugins (Frameshift, Wildtype) and population allele frequencies (--af, --af_1kg) were incorporated, along with SIFT predictions (--sift b) ^66^. Post-annotation filtering removed transcript-reference mismatches using VAtools ref-transcript-mismatch-reporter with strict mode.

### HLA Typing

HLA genotyping was performed using HLA-HD (v1.7.0) ^67^. For human Class I alleles, both classical (HLA-A, HLA-B, HLA-C) and non-classical loci (HLA-E, HLA-F, and HLA-G) were included. For Class II, the following loci were analyzed: DRA, DRB, DQA, DQB, DPA, and DPB. The HLA alleles were extracted from the HLA-HD results file (*_final.result.txt)*. Quality-controlled FASTQ files (post-Fastp processing) were aligned to the HLA reference database using Bowtie2 (v2.2.8) ^68^with multithreading (-p) and the reference index (-x) option to generate SAM files. The aligned reads were extracted using SAMtools (view -h -F 4), then converted to paired FASTQ format. Read IDs were standardized using AWK scripts, replacing the /1 and /2 suffixes with 1 and 2 for compatibility with downstream tools. HLA haplotypes were determined using *hlahd.sh* in HLAHD, with parameters including a minimum read length of 50 bp (-m 50), population allele frequency data (-f), and the HLA gene database and dictionary files.

### Host Sequences Removal

To remove host-derived genomic content, a two-step alignment strategy was implemented using the GRCh38 and telomere-to-telomere (T2T-CHM13, CHM13v2.0) ^69^human reference genomes. Quality-controlled sequencing data (after Fastp v0.22.0 preprocessing) were first aligned to GRCh38 using BWA-MEM. Unmapped reads were extracted using SAMtools, converted into FASTQ format, and subsequently realigned to the CHM13v2.0 genome. Reads that remained unmapped after both alignments were classified as putative non-host sequences. Alignment metrics including total input reads, mapped/unmapped read counts, and paired-end statistics were recorded at each step with SAMtools. For paired-end data, unmapped reads were split into R1 and R2 FASTQ files to preserve pairing information; for single-end data, unmapped reads were combined into a single output FASTQ file for each sample.

### Microbial Alignment and Taxonomic Profiling

Taxonomic classification of host-filtered reads was performed using the GATK PathSeqPipelineSpark module against a custom-curated microbial reference database. Alignments were conducted using BWA, retaining both primary and suboptimal alignments. Only reads with a minimum clipped alignment length of 50 bp were considered. To ensure taxonomic specificity, we applied a two-stage filtering strategy. For the primary alignment of each read, only mappings with alignment length ≥95% of the read length were retained. For secondary/suboptimal alignments (reported in the XA tag), both an alignment length ≥95% and a mismatch count ≤1 were required. Taxonomic identities were resolved using a custom taxonomy hierarchy file mapping genome accession to NCBI Taxonomy ID. For cross-tissue compositional comparisons, we established stringent detection thresholds to distinguish resident microbes from transient contaminants or sporadic detections. A microbial taxon (genus or species) was classified as “present in” a given tissue type (tumor or peritumor) when it was detected in ≥50% of samples from that tissue type within each cancer cohort.

### Microbial Gene Identification

To characterize microbial antigenic potential, taxonomically resolved microbial reads were aligned against our custom protein reference databset using BLASTX (v2.12.0). This in silico translation approach enables nucleotide reads to be matched against protein sequences. Only alignments with 100% identity (percentage of identical matches, Pident = 100) were retained to ensure maximal confidence in sequence assignment. BLASTX was run with the following parameters: maximum E-value of 5e-2 (-evalue 5e-2), up to five top-scoring matches per query (-max_target_seqs 5), and parallelized execution across available CPU threads (-num_threads). Output fields included query ID, subject protein ID, aligned nucleotide and protein sequences, protein annotation, percentage of identical matches, alignment length, mismatch count, gap openings, E-value, and bit score. Identified protein sequences were exported in FASTA format for downstream peptide prediction. Taxonomic identity of the matched proteins was validated via NCBI Taxonomy ID cross-referencing. Through this protein-based detection approach, we identified microbial genes expressed in each sample. A microbial gene was considered expressed if at least one read mapped to its corresponding protein sequence with 100% identity and E-value ≤ 5e-2. For quantification, the number of expressed genes was calculated by counting unique protein IDs detected per sample, with each gene counted once regardless of read depth. For tumor-specificity analysis, a gene considerd “exclusively expressed in tumor tissues” when detected with expression in tumor samples but absent in all corresponding peritumor samples from the same cancer type.

### Microbial Peptide Identification

To identify candidate antigens for HLA binding prediction, microbial protein sequences were segmented into overlapping peptides using a sliding window approach by pVACtools (v4.2.1) ^70–72^. For MHC Class I prediction, peptides of 8–10 amino acids in length were generated; for Class II, peptides of 15 amino acids were extracted. This ensured coverage of both canonical Class I and II epitope lengths.

### MHC Binding Prediction

MHC binding predictions were conducted using pVACtools (v4.2.1), an integrative pipeline that leverages multiple binding probability and eluted ligand prediction algorithms, including NetMHC, NetMHCpanEL, NetMHCIIpan, NetMHCIIpanEL, SMM, SMMPMBEC, SMMalign, and NNalign. This tool analyzes peptide binding to MHC molecules based on the HLA alleles identified in the HLA Typing step, with a focus on high-confidence, two-field resolution alleles (e.g., HLA-A02:01*). Microbial peptides generated from protein sequences (as described previously) were used as input for binding predictions. For each peptide, binding probability was predicted using multiple algorithms, and the lowest IC_50_ value across these predictions was retained as the final binding probability. Binding probability predictions were performed in parallel across multiple CPU threads (-t), utilizing IEDB tools installed in a dedicated directory (--iedb-install-directory). Peptide with an IC_50_ ≤ 500 nM was retained for further analysis, and alleles without typing information (e.g., Not_typed) were excluded to ensure reliability. For peptide comparisons, predicted HLA-binding peptides (IC₅₀ ≤ 500 nM) were considered “shared” when predicted in both tumor and peritumor tissues. Conversely, peptides were defined as ’tumor-specific’ when predicted in tumor samples but absent in all corresponding peritumor samples from the same cancer type.

### Saturation Analysis

Species accumulation curves were generated to determine sample saturation points across 11 cancers using incidence matrices in *incidence_raw* format. Rarefaction modeling was performed with the iNEXT (v3.0.1) under parameterization *q = 0* (species richness), *datatype* = “incidence_raw“. To estimate the rate of change in species richness with increasing sample size, we calculated the slope of the rarefaction curve. The slope was computed as the difference in species richness between two consecutive sample sizes, divided by the difference in sample sizes. Specifically, for each sample size 𝑡, the slope 𝑠 at each point was calculated using the formula:

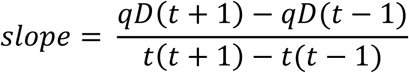

where 𝑞𝐷 represents species richness (i.e., the number of species).

The saturation point was defined as the sample size at which the rate of increase in species richness (the slope of the rarefaction curve) becomes minimal or negligible. Specifically, we calculated the percentage change in species richness (𝑞𝐷) between consecutive sample sizes. If the percentage change in species richness dropped below 5%, the corresponding sample size was considered to be the saturation point. Robust microbial saturation was achieved across all cancer samples.

### RNA-seq Coverage Calculation

Microbial reads were isolated from raw sequencing data by removing host reads. These reads were first aligned to a custom-built microbial reference database using PathSeq. The resulting BAM files were converted back to unmapped FASTQ format using SAMtools and then aligned to specific microbial reference genomes using BWA-MEM, generating a BAM file for each microorganism per sample. Coverage profiles for each sample (or merged sample group, see below) was generated from these BAM files using BEDTools (v2.31.1) ^73^ with the -bga flag.

For analyses requiring a combined coverage profile representing a specific group (e.g., pooled tumor samples, pooled peritumor samples, or defined background sample sets BG-1: 17 healthy brain, BG-2: 9 glioblastoma peritumor, BG-3: BG-1 + BG-2, BG-4: 6 healthy testis), BAM files from all samples within the group were merged using samtools merge. Coverage was subsequently calculated for these merged BAM files using BEDTools.

### Differential RNA-seq Coverage Analysis

To assess coverage differences between sample groups (e.g., tumor vs. peritumor), each microbial genome was divided into 100 non-overlapping contiguous bins. The mean RNA-seq coverage depth per bin was computed for each group and compared using a one-tailed Mann-Whitney test.

### Microbial Transcriptional Reprogramming Index

For each target sample and corresponding microorganism, two distinct genome regions were defined based on sequencing coverage: the *microbial transcriptional programming region* and the *target-specific microbial transcriptional reprogramming region*. The *microbial transcriptional programming region* corresponds to the genome regions which covered by both background and target samples, and the *target-specific microbial transcriptional **r**eprogramming (MTR) region* is defined as the region which covered by sequencing reads exclusively in target samples (e.g., tumors) but not in the background samples (Fig. 2D).

To record these regions, we constructed Boolean vector for each microorganism based on the bedGraph file, where each position (𝑣_𝑖_) in the vector represented whether sequencing reads aligned to the corresponding genome base 𝑖 (1 for aligned, 0 for not aligned). The genome-length Boolean vector 𝑉 = (𝑣_1_, 𝑣_2_, … , 𝑣_𝑛_) was created for both target and background samples.

The Microbial Transcriptional Reprogramming Index (MiTRI) was then calculated by summing the coverage values (𝐶_𝑖_) of each base 𝑖 in the *target-specific MTR region* across all 𝑛 bases, followed by normalizing with the microbial reads proportion (defined as 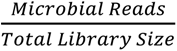), and the length of the *microbial transcriptional programming region*.

The formula for this calculation is given by:

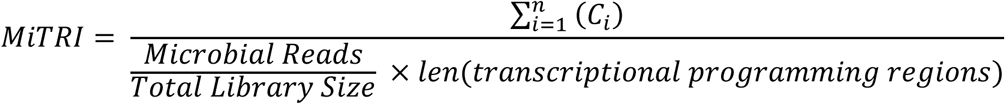

### Transcriptional Enrichment Analysis

To assess whether MTR represents selective transcriptional activation rather than abundance-driven effects, we calculated transcriptional enrichment scores for tumor-associated bacteria across all tumor samples. For each bacterium 𝑖 in detected in tumor sample 𝑗 , we identified tumor-specific MTR regions using peritumor tissues of the corresponding cancer type as background. The enrichment score was defined as:

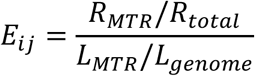

where 𝑅_𝑀𝑇𝑅_means number of reads mapping to tumor-specific MTR regions, 𝑅_𝑡𝑜𝑡𝑎𝑙_ means total reads mapping to the bacterial genome, 𝐿_𝑀𝑇𝑅_ means sum of all tumor-specific MTR region lengths, and 𝐿_𝑔𝑒𝑛𝑜𝑚𝑒_complete genome length of the bacterium. This ratio compares the observed read density in MTR regions to the expected density under uniform distribution; an enrichment score of 1 indicates reads are uniformly distributed across the genome as expected by chance, while scores > 1 indicate preferential transcription of MTR regions beyond their genomic representation.

Statistical significance was evaluated using a two-sided binomial test with the null hypothesis that reads are uniformly distributed across the genome. Sample-bacteria pairs (each representing one bacterium in one tumor sample) with *p* < 0.05 were considered to show significant differential transcription and included in enrichment analysis. Enrichment scores were log₂-transformed for visualization.

### Gene Ontology (GO) Enrichment Analysis

Enrichment analysis of microbial peptides was performed by analyzing the source predicted microbial protein-coding sequences that were expressed exclusively in either tumor or peritumor tissues. To further investigate the GO functional enrichment of the proteins, we first collected annotations for all samples from our custom annotated protein database. GO annotations of the proteins were retrieved from the UniProt (https://www.uniprot.org/id-mapping) using sequence identifiers from our custom protein database. Due to the query limit of 100,000 identifiers per submission on the UniProt website, the 10,483,367 sequence identifiers in our protein database were divided into 105 batches for annotation retrieval, resulting in 5,182,522 records of 2,878,757 protein sequence identifiers.

For each GO term 𝑔, a hypergeometric test was used to assess its enrichment in tumor and peritumor groups. The following statistical parameters were computed:

𝑁: Total number of proteins annotated with at least one GO term.

𝑛: Number of proteins specifically annotated with GO term 𝑔.

𝑀: Number of identified proteins in the group annotated with at least one GO term.

𝑚: Number of identified proteins in the group annotated with GO term 𝑔. The *P* value was calculated using the hypergeometric test:

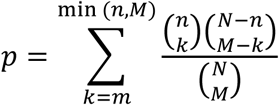

where 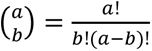 represents the binomial coefficient, indicating the number of ways to choose 𝑏 elements from 𝑎. Finally, GO terms were considered significantly enriched in tumor or peritumor samples if their adjusted *P* value was less than 0.05 corrected by Benjamini-Hochberg (FDR-BH).

### Microbial Abundance Quantification

Microbial abundance were quantified as normalized relative eRPKM, calculated as:

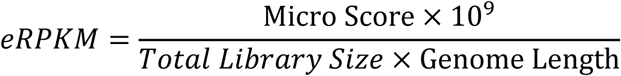

where *micro score* represents the number of reads mapped to a given taxon, weighted by 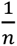 for reads aligned to 𝑛 reference genomes (i.e., multi-mapped reads). Multi-mapped reads were resolved by proportionally distributing their contributions 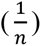 across all valid alignment targets. The resulting values were then scaled to percentages, representing the relative abundance of each microbial taxon in each sample.

### Jaccard Similarity Indexes Calculation

Somatic mutations were extracted from VCF files by retaining variants with “PASS” status in the *FILTER* column. Each mutation was annotated with its genomic coordinates (chromosome, start position) and alteration type (single-nucleotide variant, insertion, or deletion). Unique mutation identifiers were created by concatenating the chromosome, start position, and alteration type (e.g., “chr1_123456_A>T“). Pairwise Jaccard indices between samples were computed as the ratio of shared mutated driver genes (intersection) to the total distinct mutated driver genes (union) across all sample pairs, generating a similarity matrix for downstream analysis. The Jaccard similarity for two sets of genes 𝐴 and 𝐵 is given by:

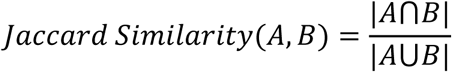

where |𝐴⋂𝐵| is the number of shared mutated driver genes between the two samples and |𝐴⋃𝐵| is the total number of distinct mutated driver genes across both sets.

### Enrichment Analysis of Differentially Expressed Host Genes

Differential gene expression analysis was performed using DESeq2 (v1.42.1) ^74^ on raw RNA-seq count matrices. Count data were filtered to remove duplicate gene symbols, retaining unique entries for further analysis. The DESeq function applied default normalization and dispersion estimation, followed by Wald testing for differential expression. For enrichment analysis, differentially expressed genes (absolute log2-fold change ≥ 1 and *p* < 0.05) were ranked by their log2-fold change values. Gene symbols were mapped to ENTREZ IDs using the org.Hs.eg.db annotation database. Biological pathway enrichment for GO terms was conducted using clusterProfiler (v4.10.1) ^75^ with Benjamini-Hochberg FDR correction (adjusted p <0.05), focusing on biological process (BP) categories. The enrichment results were visualized using dot plots displaying the top 25 significant terms, with point sizes scaled to gene ratios and colors indicating FDR-adjusted *p* values.

### Deconvolution Analysis of bulk RNA-seq data

To estimate the relative abundance of immune cell populations in bulk RNA-seq samples, we performed gene expression deconvolution using CIBERSORT (v0.1.0) ^76^with the LM22 signature matrix. Prior to deconvolution, genes were filtered based on expression levels to reduce noise, retaining only those with TPM ≥ 1 in at least 50% of the samples. CIBERSORT was then executed with 1000 permutations and without quantile normalization to ensure consistency across datasets.

### Preprocessing of single-cell RNA-seq data

Low-quality reads were filtered from single-cell RNA sequencing (scRNA-seq) raw data using fastp (v0.23.2) with default parameters. High-quality reads were aligned to the GRCh38 human reference genome using STAR (v2.7.10a). Raw gene expression matrices were quality-controlled by retaining cells expressing at least 200 genes and genes detected in a minimum of 3 cells. After removing low-quality cells, potential doublets were identified using DoubletDetection (v4.2)^77^. The final dataset comprised 43,502 high-confidence single cells.

The Scanpy (v1.10.2) was used for data scaling, transformation, clustering, dimensionality reduction, differential expression analysis and most visualization. Counts were normalized to 10,000 transcripts per cell using total-count normalization, followed by log1p transformation. Highly variable genes (HVGs) were selected based on mean expression (0.01 < mean < 5) and dispersion (dispersion > 0.3), Retaining the top 3,000 highly variable genes and marker genes (e.g., CD3D, CD4, CD8A) for downstream analysis. Expression values were scaled to unit variance (clipped at a maximum value of 10) and subjected to principal component analysis (PCA) using the ARPACK solver (50 components). A KNN graph was constructed using the first 10 principal components (15 neighbors per cell). The PCA subspace was further visualized by UMAP. Batch effects across samples were corrected using BBKNN (Batch Balanced k-Nearest Neighbors).

### T Cell Subset Annotation

Firstly, T cells were identified by CD3D and CD3E and annotated by classical marker genes for CD4^+^, CD8^+^, and γδ T cells (CD4 T: CD4; CD8 T: CD8A and CD8B; γδ T: TRGV9 and TRDV2). Then, an additional round of preprocessing steps was performed to classified the subtype of CD4^+^ and CD8^+^T cells with a resolution of 1.2 and 0.8 respectively. Finally, each type of CD4 T cells (naive/central memory T cells, Tn/Tcm; effector memory RA/effector T cells, Temra/Teff; regulatory T cells, Treg) and CD8 T cells (naive/central memory T cells, Tn/Tcm; effector memory T cells, Tem; effector memory RA/effector T cells, Temra/Teff) were annotated using classical marker genes (Extended Data Fig. S22; Extended Data Table S7).

### Single-cell TCR data Analysis and TCR-Peptide Affinity Prediction

Quality control of raw scTCR sequencing data was performed with fastp (v0.23.2). Then, Cell Ranger VDJ (v7.2.0) was conducted to identify TCR α- and β-chain sequences and generate the clonotype information for each cell with default parameters. Clone abundance of each clonotype were calculated, and clonotypes observed more than once were considered clonally expanded. For clones with completed α and β-chain information, Net-TCR (v2.2) was employed to predict the binding affinity between TCR clonotypes and candidate epitope peptides^78^. Predicted binding affinity scores were categorized as follows: high affinity (≥ 0.6), intermediate affinity (≥ 0.4 and < 0.6), and low affinity (<0.4).

### Structural Modeling and Dynamics Simulation of TCR-pMHC Interaction

To elucidate the structural basis of TCR recognition of the peptide–MHC (pMHC) complex, we employed a multi-step computational pipeline. The initial crystal structure of the MHC molecule (e.g., HLA-C*08:02) was obtained from the Protein Data Bank (PDB), and non-crystallographic water molecules were removed using PyMOL^79^. The co-crystallized peptide was separated, and the target peptide sequence was modeled by mutating the original peptide using PyMOL’s Mutagenesis Wizard. The modeled peptide was then redocked into the MHC binding groove using HDOCK (v1.1) ^80^, and the top 10 ranked pMHC complexes were evaluated by structural inspection in PyMOL. The most structurally plausible model—based on docking score and binding geometry—was selected for downstream TCR docking. For the TCR, the variable regions of both α and β chains were concatenated and submitted to AlphaFold3 for structure prediction ^81^. The model with the highest pLDDT score was retained and used as the receptor for docking. The selected TCR model was docked onto the pMHC complex (as ligand) using HDOCK. Among the 10 predicted docking poses, the final TCR–pMHC complex was manually selected based on docking rank and the proximity of the TCR CDR3β loop to the pMHC interface.

Molecular dynamics (MD) simulations were performed using GROMACS (v2024.3) with the AMBER99SB-ILDN force field ^82^. The ternary complex was solvated in a cubic box using the SPC/E water model with at least 1.5 nm padding, and neutralized with Na⁺ or Cl⁻ ions. Energy minimization was performed via steepest descent until the maximum force was below 1000.0 kJ/mol/nm. Equilibration was conducted in two steps: a 1 ns NVT ensemble using a V-rescale thermostat at 300 K with position restraints on protein heavy atoms, followed by a 1 ns NPT ensemble using a Parrinello-Rahman barostat at 1.0 bar. Production MD was run for 100 ns with all restraints released. LINCS constraints were applied to bonds involving hydrogen, allowing a 2 fs timestep; PME was used for long-range electrostatics, and a cutoff of 1.2 nm was set for both Coulomb and van der Waals interactions.

Binding free energy (ΔG) between the TCR and pMHC was estimated using the MM/PBSA method via gmx_MMPBSA (v1.6.4) ^83^. Snapshots from the equilibrated phase of the MD trajectory were preprocessed using gmx trjconv with -pbc mol and -center to remove periodic boundary artifacts. The processed trajectory was analyzed under the Generalized Born model (igb=5) to compute ΔG values.

### High Performance Computing (HPC) Based Data Processing

Numerical computations were performed on the AMD EPYC 7702-based cluster at the Hefei Advanced Computing Center. Key hardware specifications include:

CPU: Dual AMD EPYC 7702 processors (128 physical cores / 256 threads per node)

Memory: 256 GB DDR4 RAM per node

Interconnect: 100 Gb/s Omni-Path Architecture (OPA)

The microbial antigen analysis involved the following steps and estimated core-hour requirements per sample:

Quality Control (QC): 10 core-hours Expression Quantification: 30 core-hours

HLA Typing: 200 core-hours

Alignment to Reference Genomes: 40 core-hours Microbial Quantification: 20 core-hours

Protein Database Matching: 100 core-hours MHC Binding Affinity Prediction: 50 core-hours

All computational time estimates represent averaged values. Actual durations varied depending on factors such as sequencing depth and real-time computational load. In total, approximately 750,000 core-hours and 1,000 GPU-hours were consumed to analyze 1,868 pan-cancer bulk RNA-seq public samples, along with in-house generated WTS, WES, and scRNA datasets. Specific steps for 134 public bulk RNA-seq samples exceeded average runtime thresholds by orders of magnitude. Consequently, 1,734 public samples were successfully analyzed.

Structural modeling and molecular dynamics (MD) simulations for the TCR-pMHC interaction were carried out on a high-performance computing platform provided by Virtaitech (Shanghai) Co., Ltd. Each compute instance was configured with:

CPU: Intel® Xeon® Gold 6342 @ 2.80GHz, 24 physical cores; GPU: 2 × NVIDIA RTX 3090 GPUs, each with 24 GB of memory.

In total, 5 compute nodes (10 GPUs) were employed for parallel execution.

## Data availability

The microbial reference database used in this study, including reference genomes and protein sequences downloaded from the National Center for Biotechnology Information (NCBI), is publicly available at Zenodo (https://doi.org/10.5281/zenodo.15043056). Gene Ontology (GO) annotations were obtained from UniProt (https://www.uniprot.org/).

Public whole-transcriptome sequencing (WTS) data for pan-cancer samples were downloaded from the Gene Expression Omnibus (GEO). The GEO project accession numbers for each cancer type are provided in Supplementary Extended Data Table S2. Processed pan-cancer analysis results are available at Zenodo (https://doi.org/10.5281/zenodo.15041208 and https://doi.org/10.5281/zenodo.15043056). Sequencing data generated in this study have been deposited in the Sequence Read Archive (SRA) under BioProject accession number PRJNA1301281 (controlled access; requests for access should be directed to the corresponding author).

## Code availability

All analysis code is publicly available at GitHub (https://github.com/BioStaCs-public/MiTRI/). Additional supplementary data are provided with this paper.

## Acknowledgments

We sincerely express our gratitude to the Gastrointestinal oncology Integrated Research Team of Hefei Cancer Hospital of CAS for their assistance in sample extraction, data organization, standardization of clinical cases and pathological prognosis evaluation. We gratefully acknowledge the Hefei Advanced Computing Center for providing computational resources and technical support.

We are deeply indebted to Professor Qian Zhao for her expert guidance on microbial epitope discovery methodologies. Her pioneering work in low-abundance antigen detection provided critical intellectual foundation for this study.

We thank the LSCCB under health@InnoHK Program launched by ITC, The Government of Hong Kong SAR, and also Prof Chi-Ming Che for generous support on instrument.

We thank the Research Grant Council (RGC) Hong Kong, Research Committee of Hong Kong and Hong Kong Health Bureau for their kind support of this project.

We are grateful to Professor Mingwei Li from the Department of Physics, University of Science and Technology of China, for her valuable guidance on molecular dynamics simulations.

We also thank the colleagues in our research group for inspiring discussion and their contributions.

## Author contributions

Conceptualization: Z.Y.W., Z.L., F.J.M.

Methodology: F.J.M., Z.Y.W., Z.Y.X., W.W., Z.X., C.T., Z.L., L.X.T., L.K.C., Z.Q.

Investigation: F.J.M., Z.Y.X., W.W., Z.X., C.T., L.K.C., Y.H.Y., Z.T.Y., Z.Y.L., W.D.D.,

Z.J.F., C.J.L., T.Z., Z.S.W., W.L.R., W.W., L.Z.L., D.D.S., S.R.X., Z.D.D., H.Y.,

Z.Y.F., Y.C.K., C.D.D., L.X.T., J.L., Z.Q., Z.L., W.H.Z., Z.Y.W.

Visualization: Z.X., Z.Y.X., W.W., C.T., Y.H.Y.

Funding acquisition: Z.Y.W., F.J.M., Z.L., W.H.Z.

Project administration: W.H.Z., F.J.M., Z.Y.W.

Supervision: Z.L., W.H.Z., Z.Y.W.

Writing – original draft: Z.Y.W., Z.Y.X., C.T., Z.X., W.W., F.J.M.

Writing – review & editing: W.W., Z.X., Z.Y.X., L.X.T., W.H.Z., Z.Q., Z.L., F.J.M., Z.Y.W.

---

**Extended Data Fig. S1.**
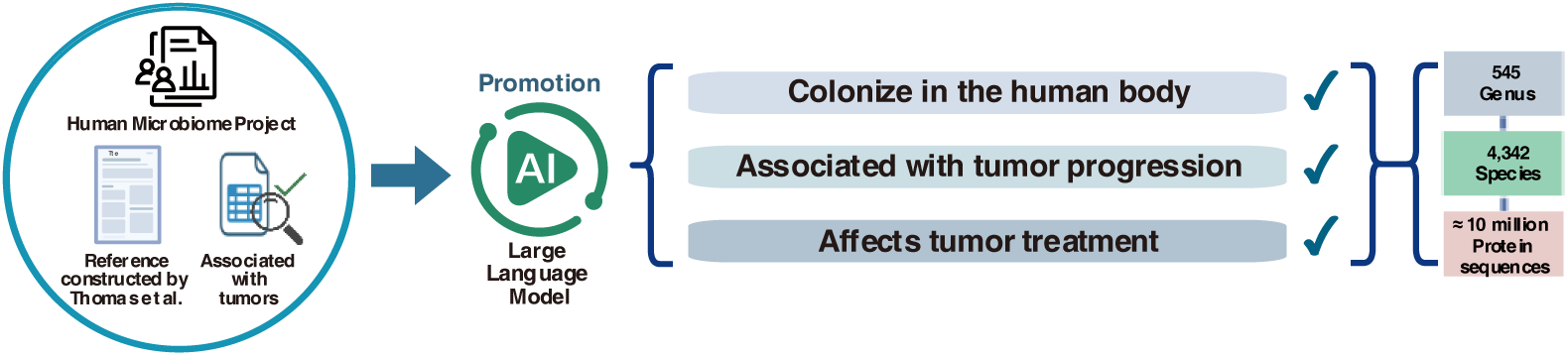
Construction of microbial reference genomes and protein sequence datasets based on Large Language Model and manual curation.

**Extended Data Fig. S2.**
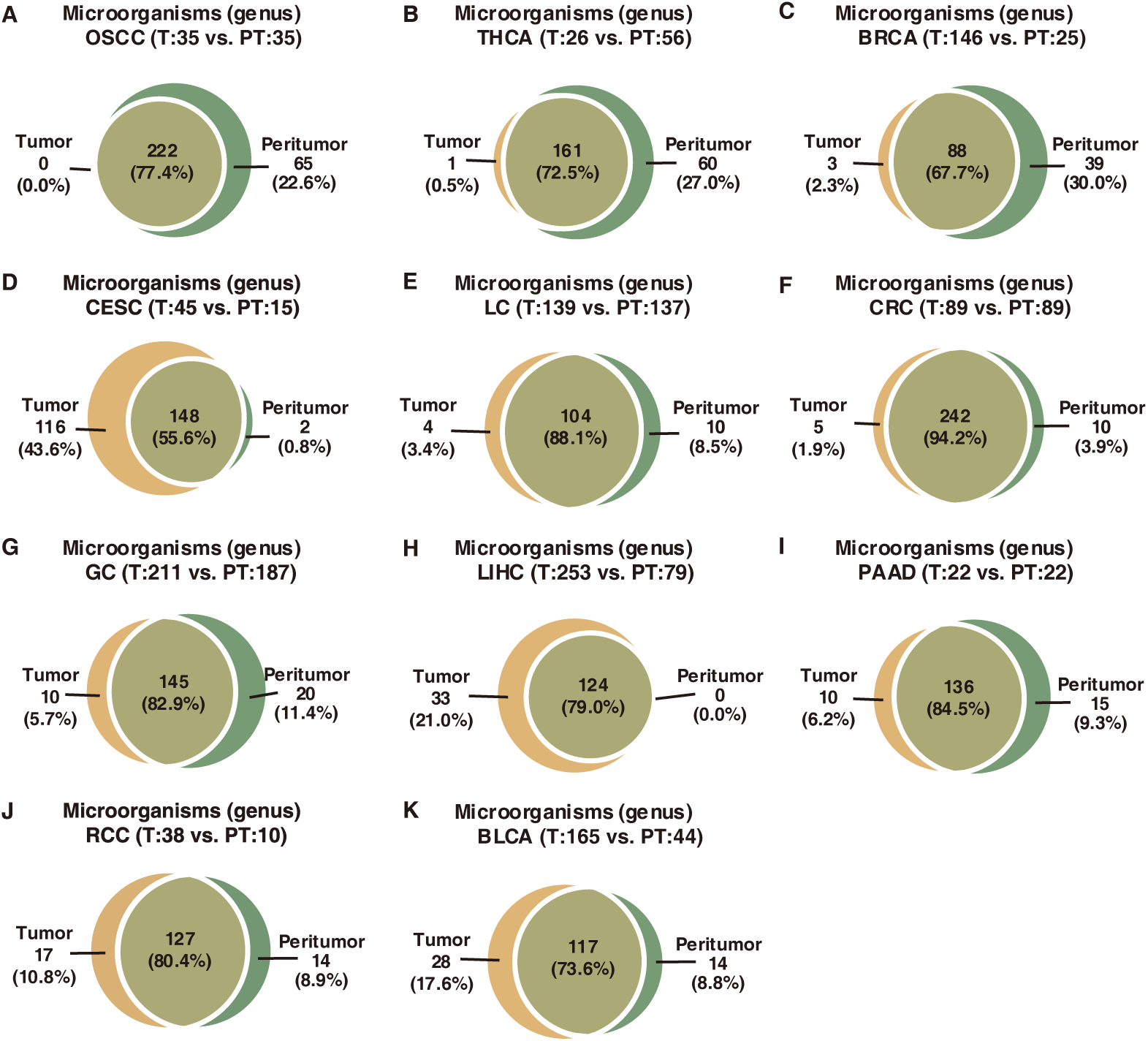
Sharing rate of microbial genus between tumor and peritumor tissue samples across 11 cancer types: bladder cancer (BLCA), breast cancer (BRCA), cervical cancer (CESC), colorectal cancer (CRC), renal cell carcinoma (RCC), liver cancer (LIHC), lung cancer (LC), oral squamous cell carcinoma (OSCC), pancreatic cancer (PAAD), gastric cancer (GC), and thyroid cancer (THCA).

**Extended Data Fig. S3.**
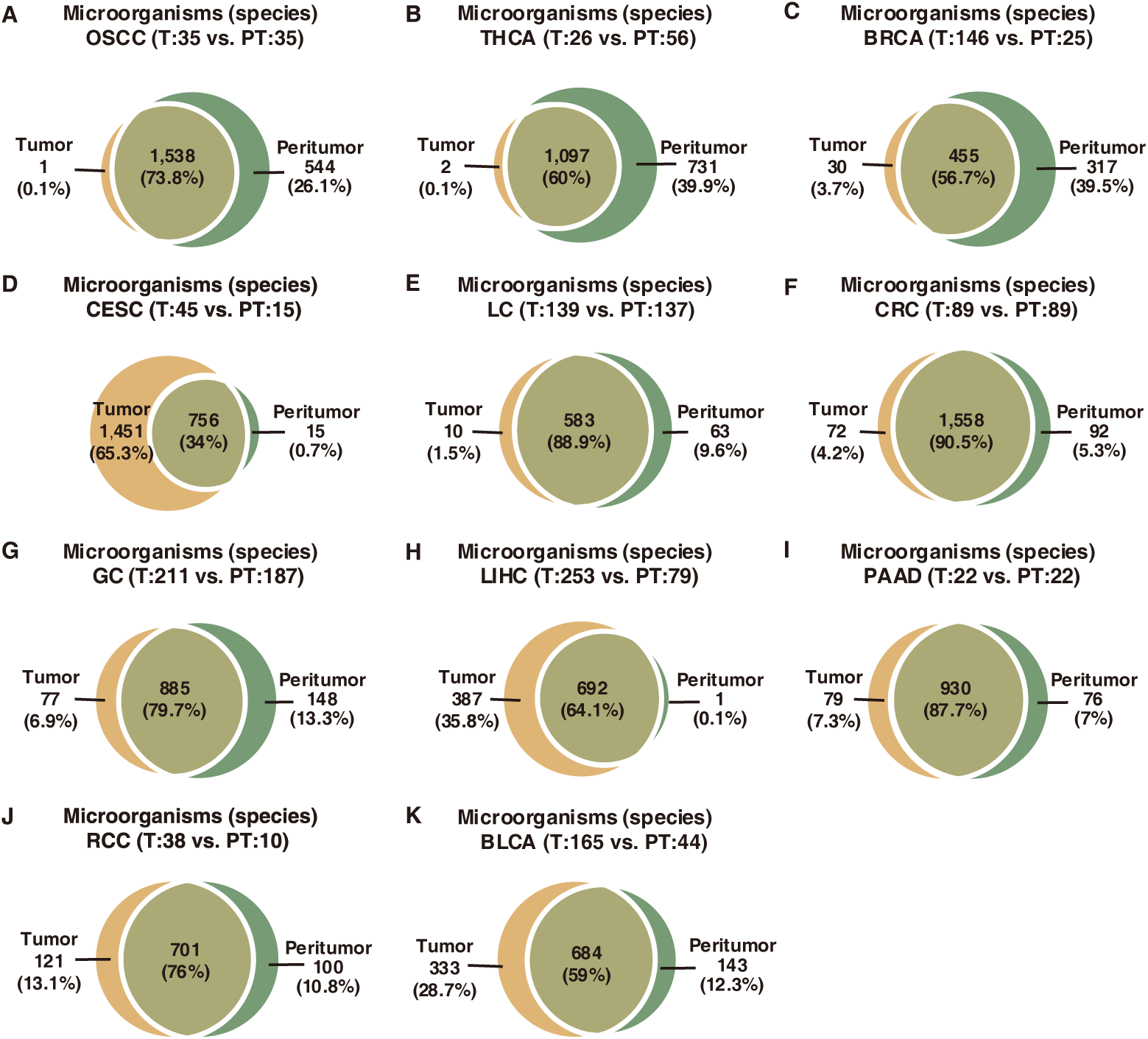
Sharing rate of microbial species between tumor and peritumor tissue samples across 11 cancer types.

**Extended Data Fig. S4.**
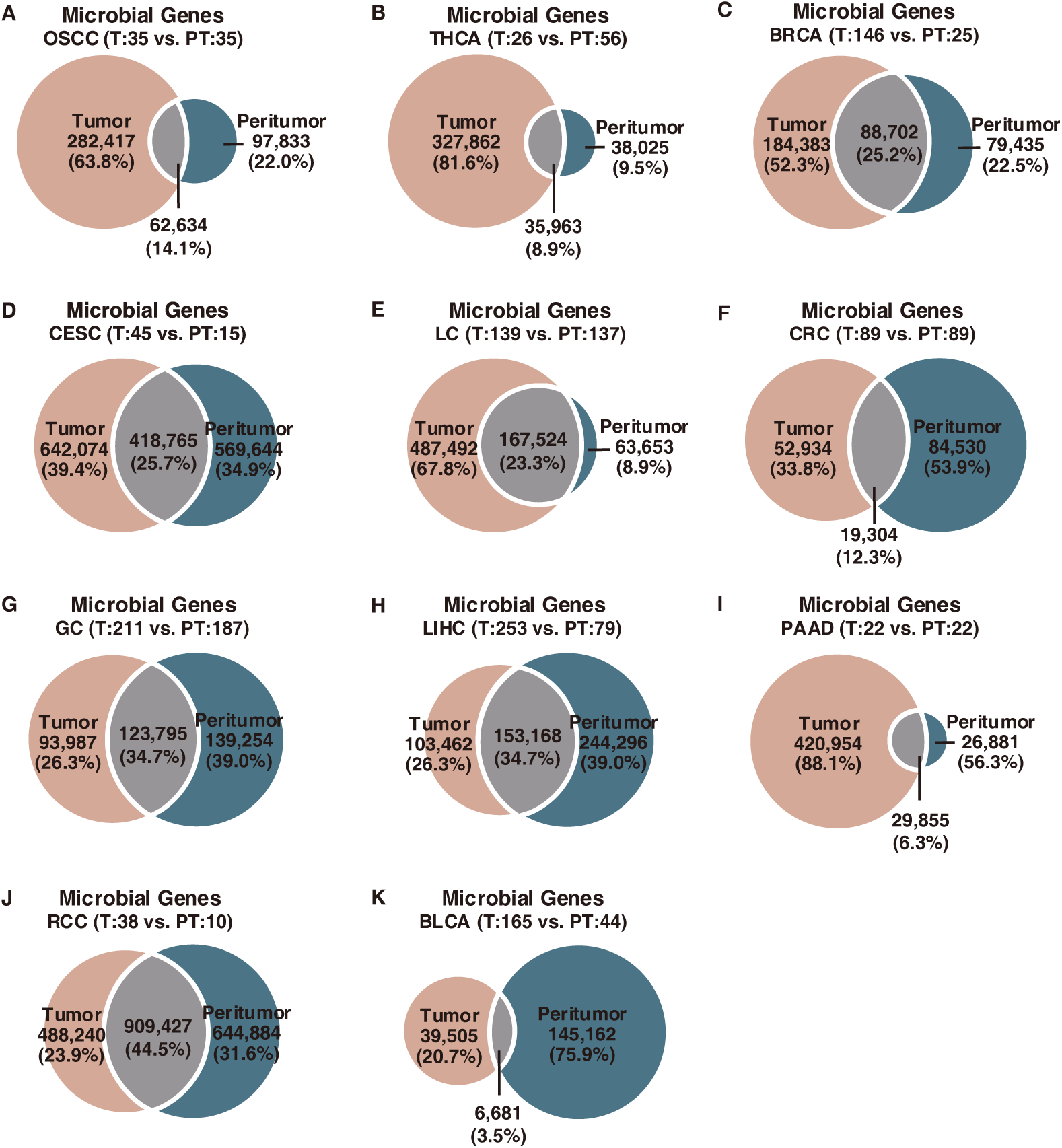
Sharing rate of microbial expressed genes between tumor and peritumor tissue samples across 11 cancer types.

**Extended Data Fig. S5.**
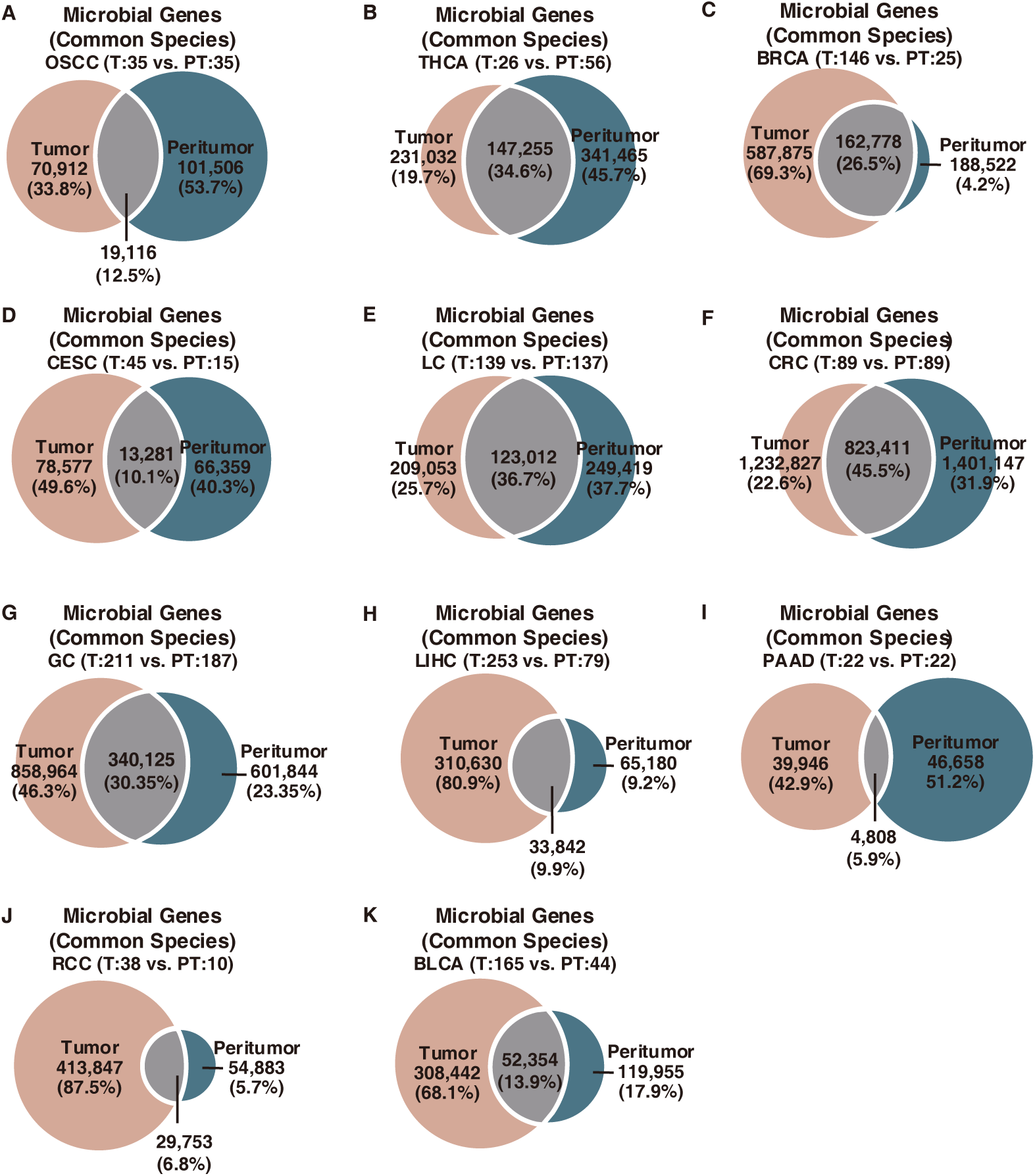
Sharing rate of expressed genes derived from shared microbial species between tumor and peritumor tissue samples across 11 cancer types.

**Extended Data Fig. S6.**
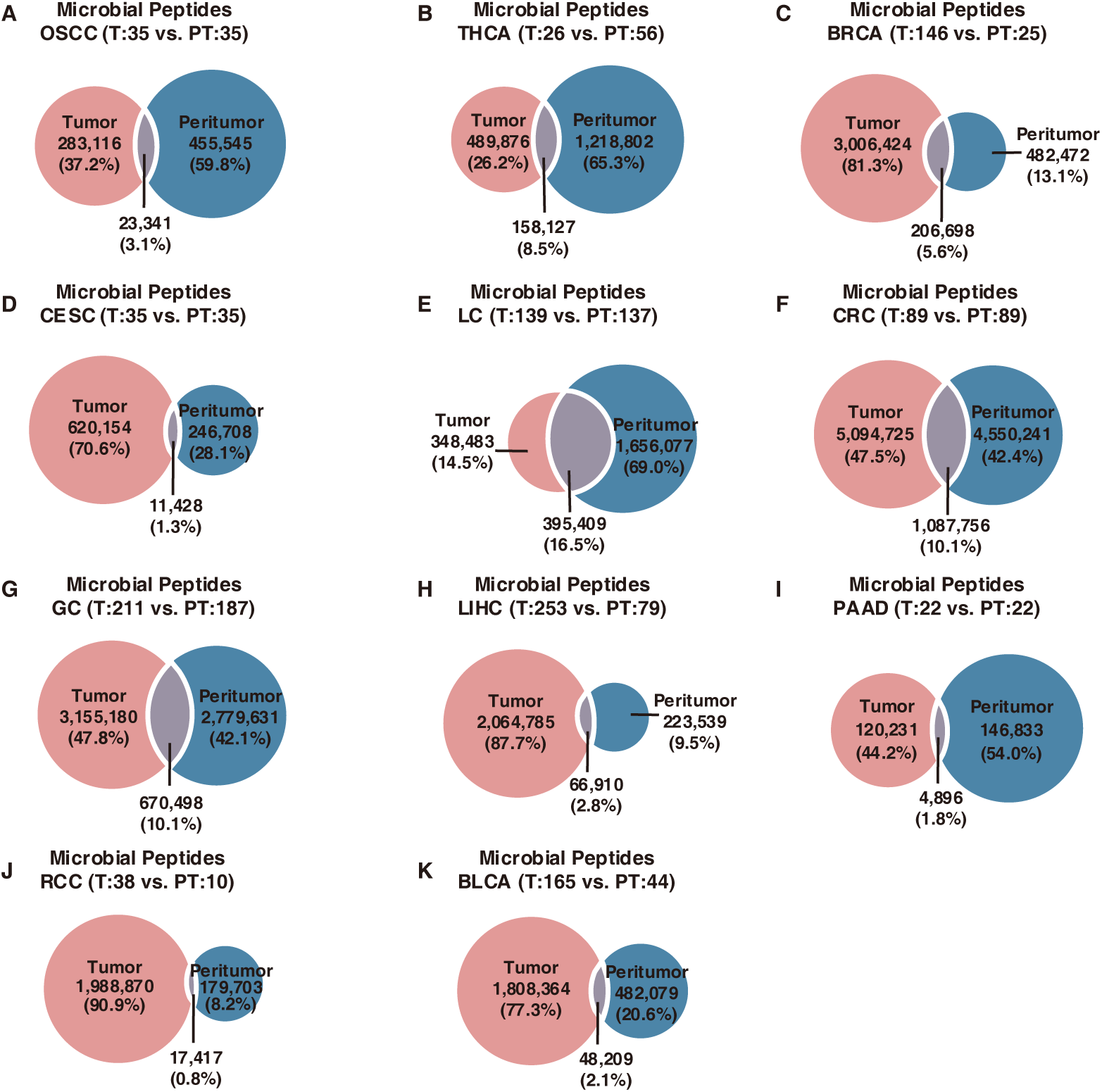
Sharing rate of predicted microbial HLA-binding peptides potential (IC_50_ ≤ 500 nM) between tumor and peritumor tissue samples across 11 cancer types.

**Extended Data Fig. S7.**
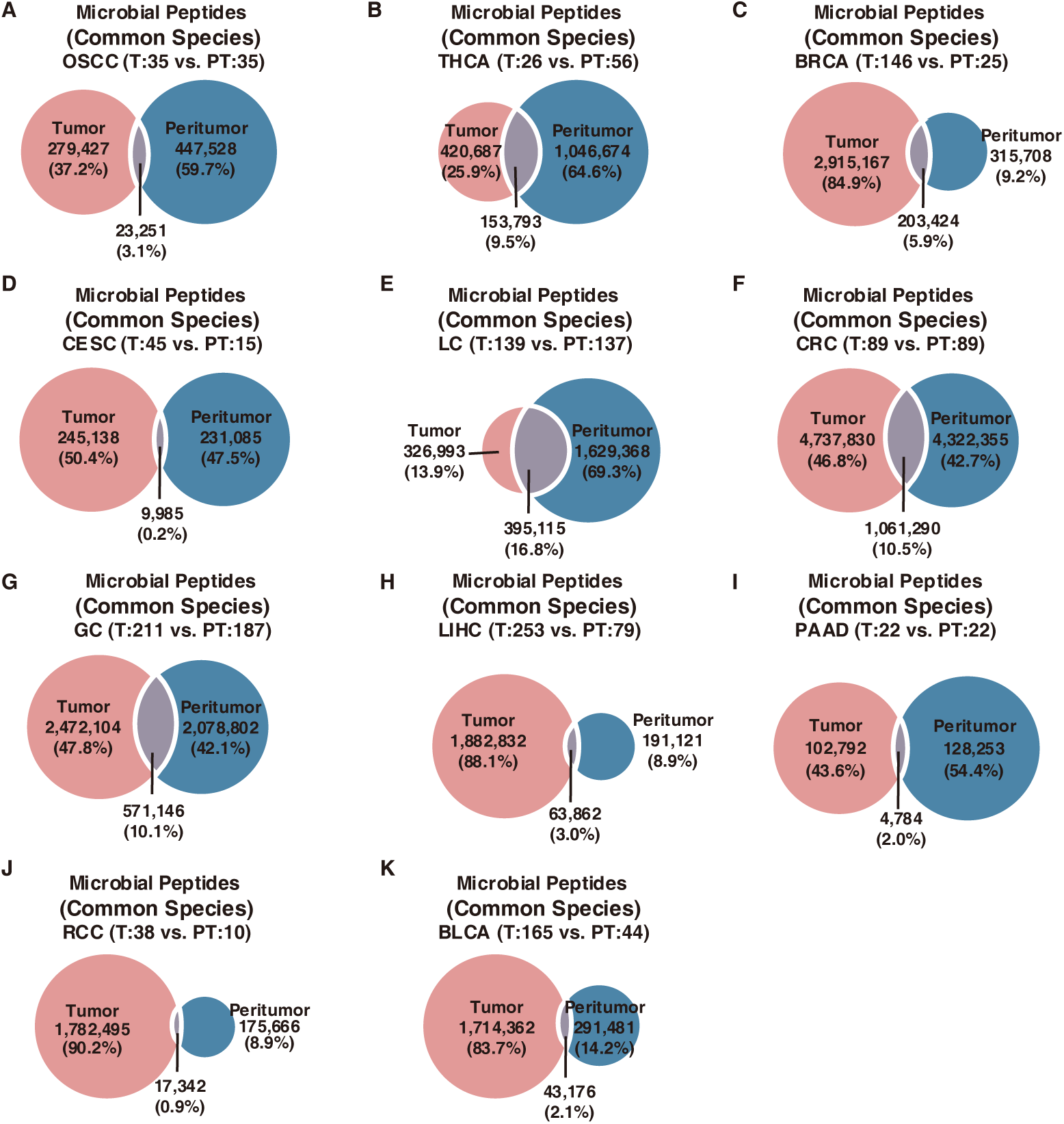
Sharing rate of predicted microbial HLA-binding peptides potential (IC_50_ ≤500 nM) derived from microbial species resident in both tumor and peritumor tissue samples across 11 cancer types.

**Extended Data Fig. S8.**
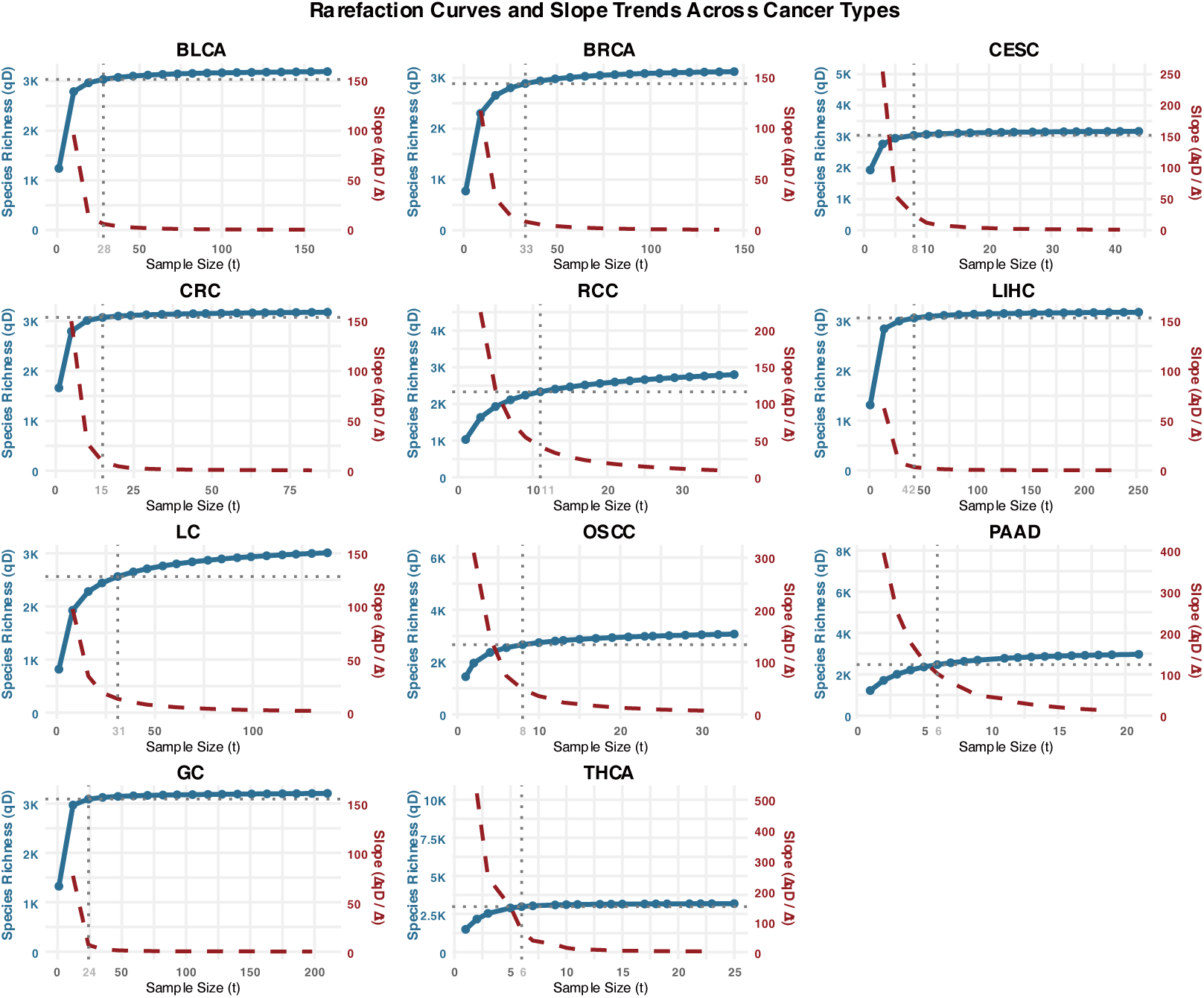
Rarefaction Curves and Slope Trends Across Cancer Types. For each cancer type, the rarefaction curves (blue lines) represent species richness (𝑞𝐷) as a function of sample size (𝑡). The dashed red lines indicate the calculated slope of the rarefaction curve, illustrating the rate of change in species richness with increasing sample size. The vertical dotted lines indicate the sample size at which the curve reaches saturation, defined as the point where the slope becomes minimal (i.e., the percentage change in species richness drops below 5%). The number next to each cancer type represents the sample size (𝑡) at which saturation occurs. All data were derived from rarefaction analysis using the iNEXT method.

**Extended Data Fig. S9.**
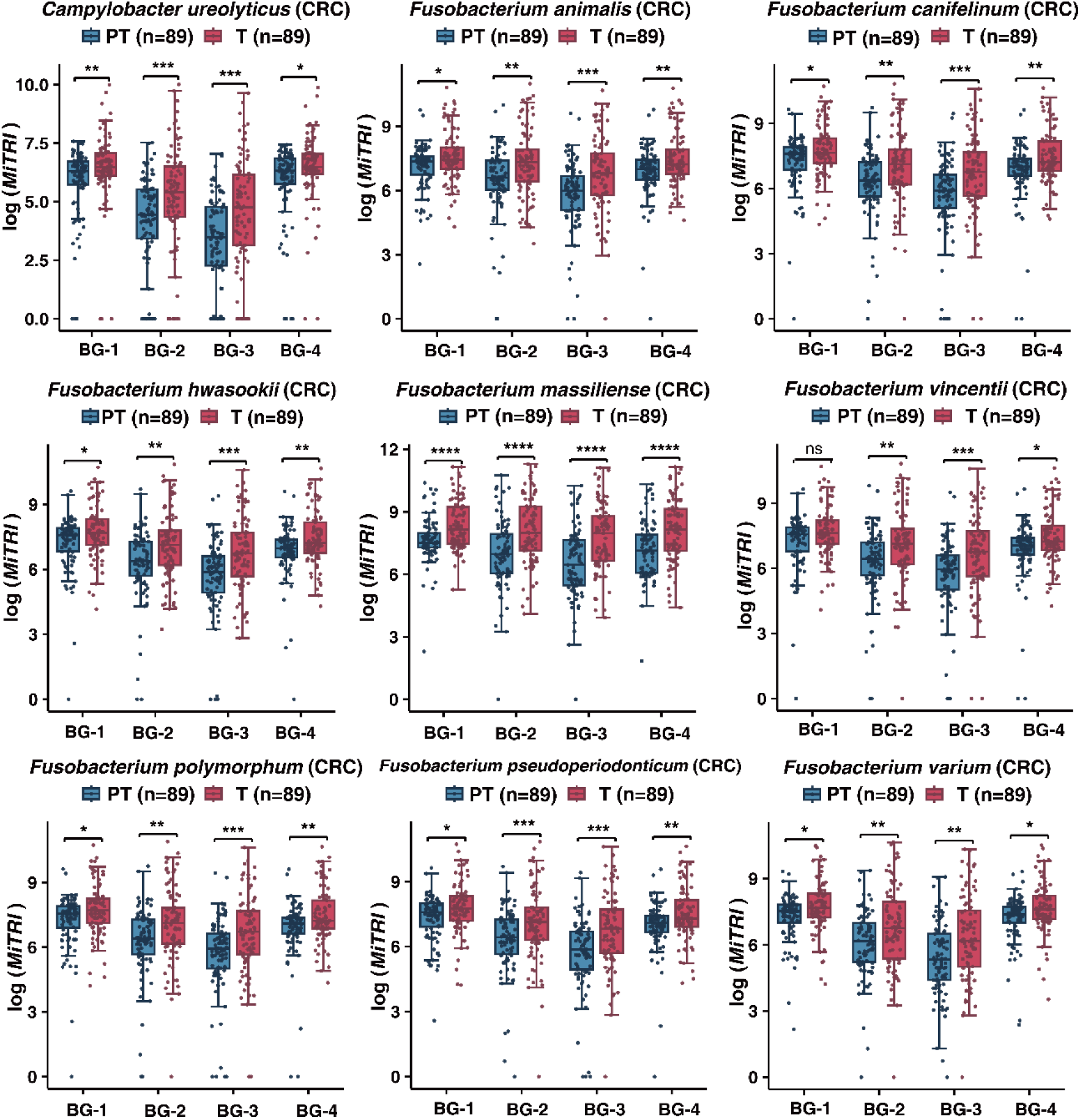
Microbial Transcriptional Reprogramming Index (MiTRI) of literature-reported tumor-associated bacteria in CRC.

**Extended Data Fig. S10.**
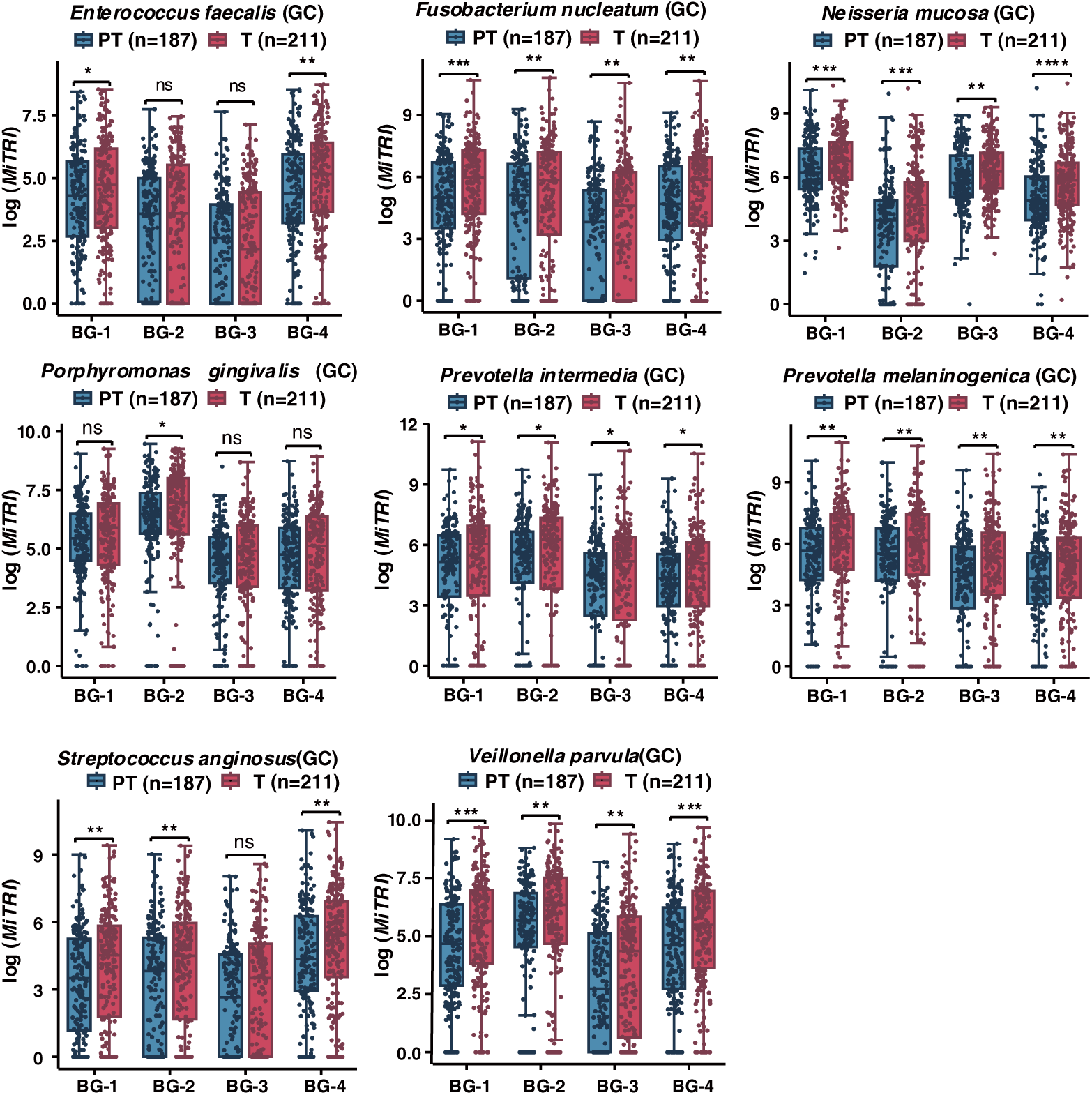
Microbial Transcriptional Reprogramming Index (MiTRI) of literature-reported tumor-associated bacteria in GC.

**Extended Data Fig. S11.**
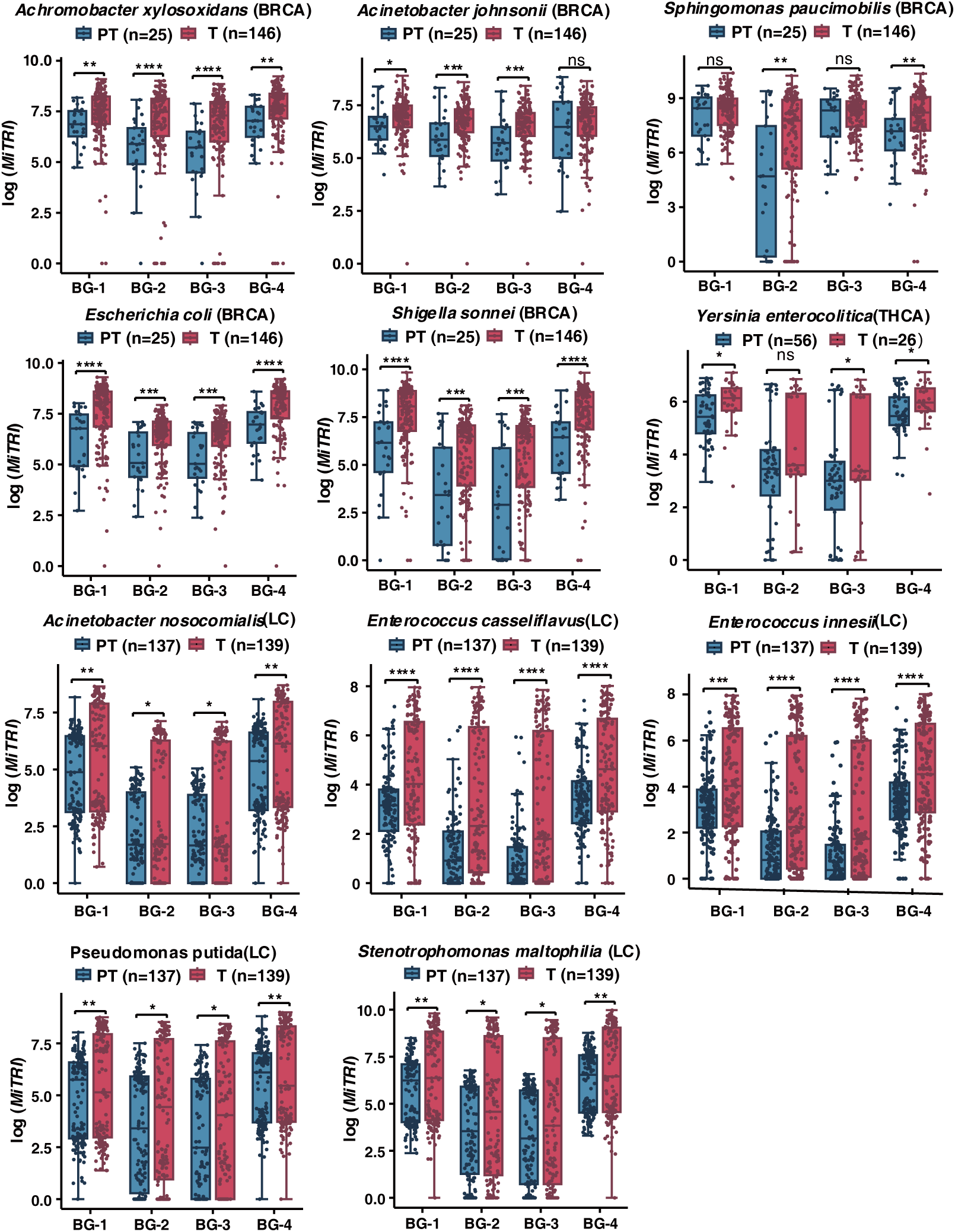
Microbial Transcriptional Reprogramming Index (MiTRI) of literature-reported tumor-associated microbes in BRCA, THCA, and LC.

**Extended Data Fig. S12.**
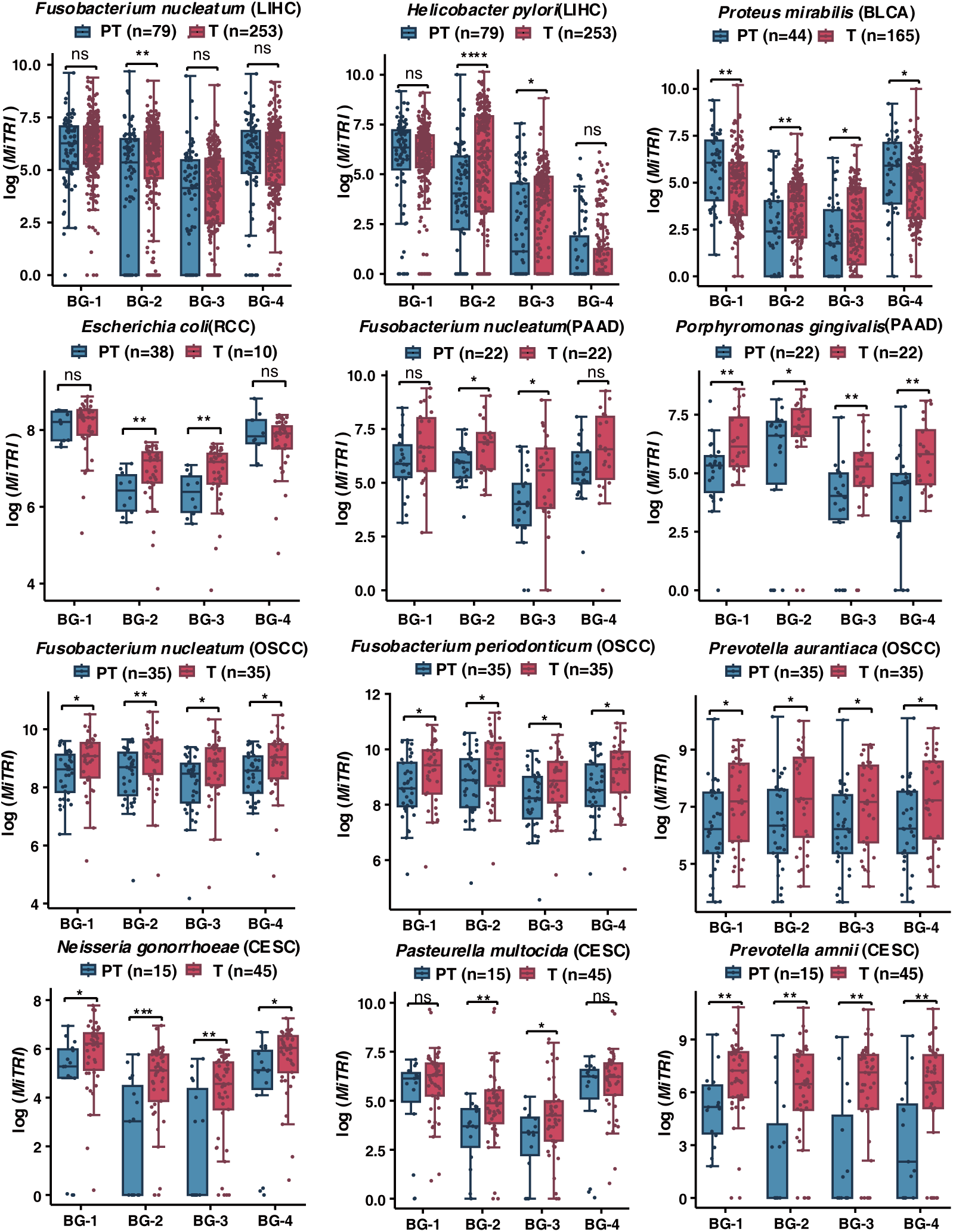
Microbial Transcriptional Reprogramming Index (MiTRI) of literature-reported tumor-associated microbes in LIHC, BLCA, RCC, PAAD, OSCC, and CESC.

**Extended Data Fig. S13.**
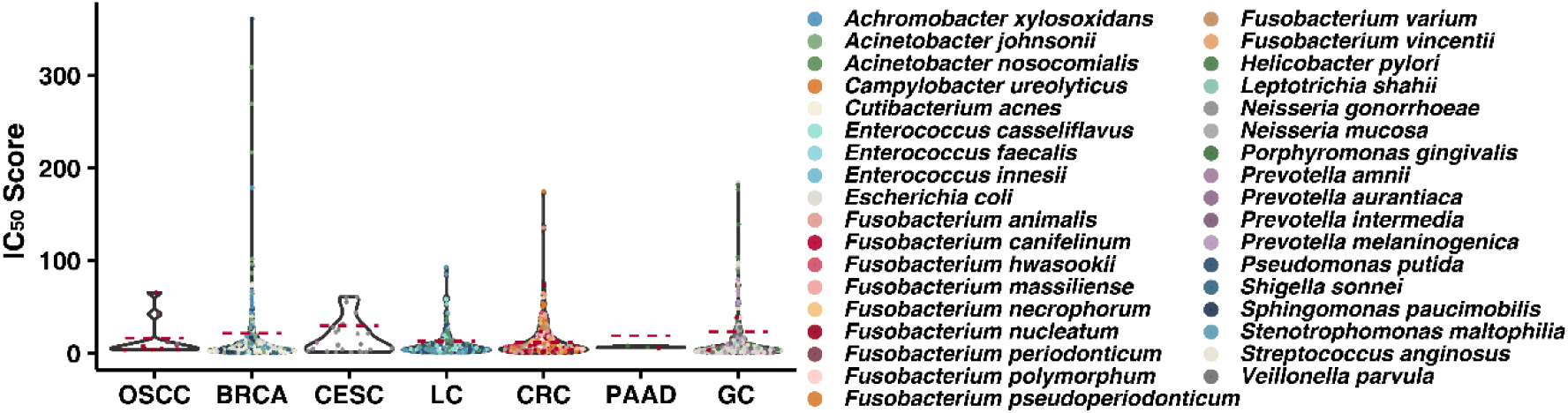
Distributions of tumor-specific MiTRI and predicted IC_50_ of tumor-specific peptides. Tumor-specific predicted microbial HLA-binding peptides derived from MTR regions exhibited stronger HLA binding probability (lower IC_50_) compared to the general predicted HLA-binding peptides derived from genome-wide regions (red dash: lowest IC_50_ of general predicted HLA-binding peptides for each microbe).

**Extended Data Fig. S14.**
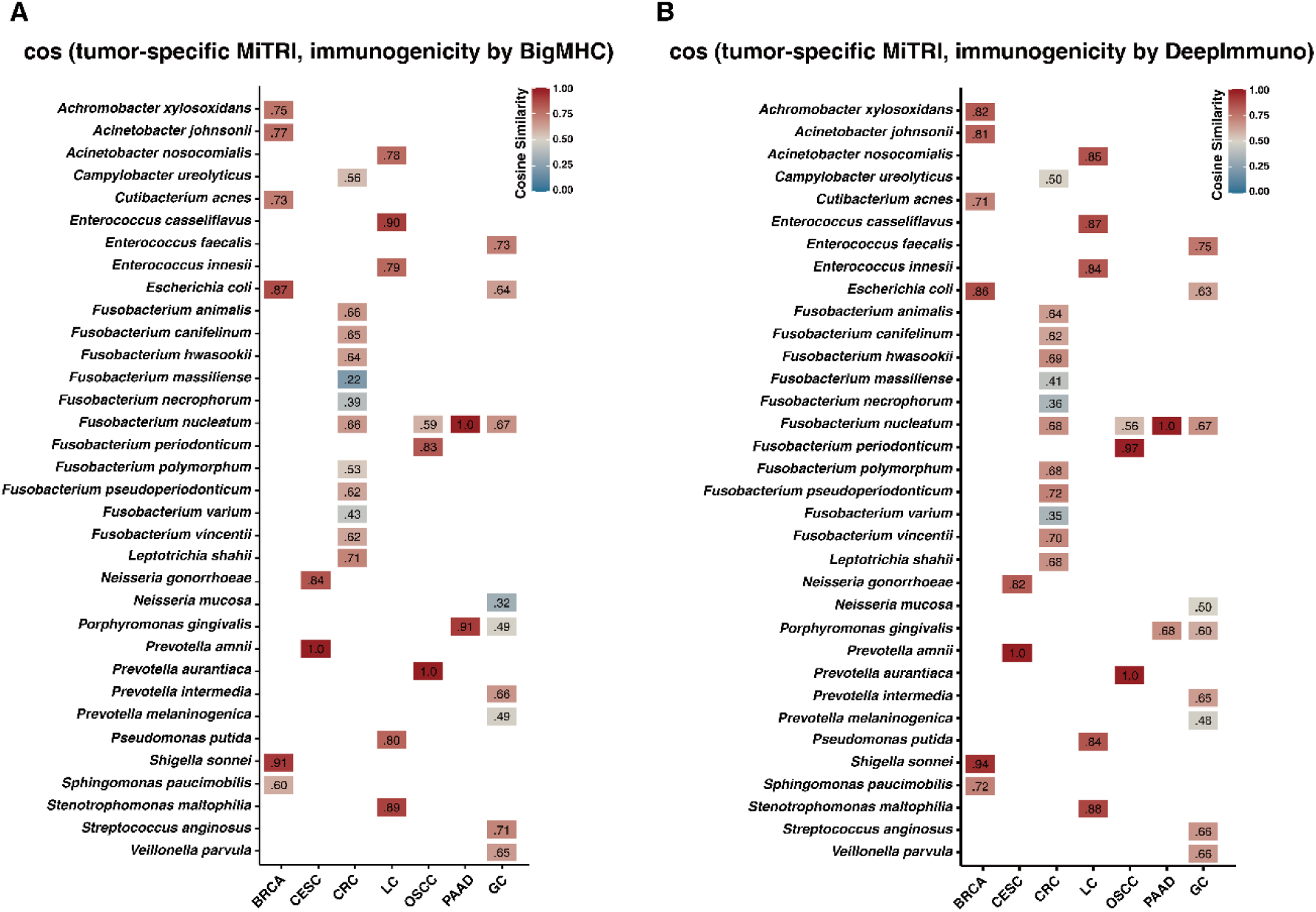
Cosine similarity of tumor-specific MiTRIs and immunogenicity predicted by **(A)** BigMHC and **(B)** DeepImmuno.

**Extended Data Fig. S15.**
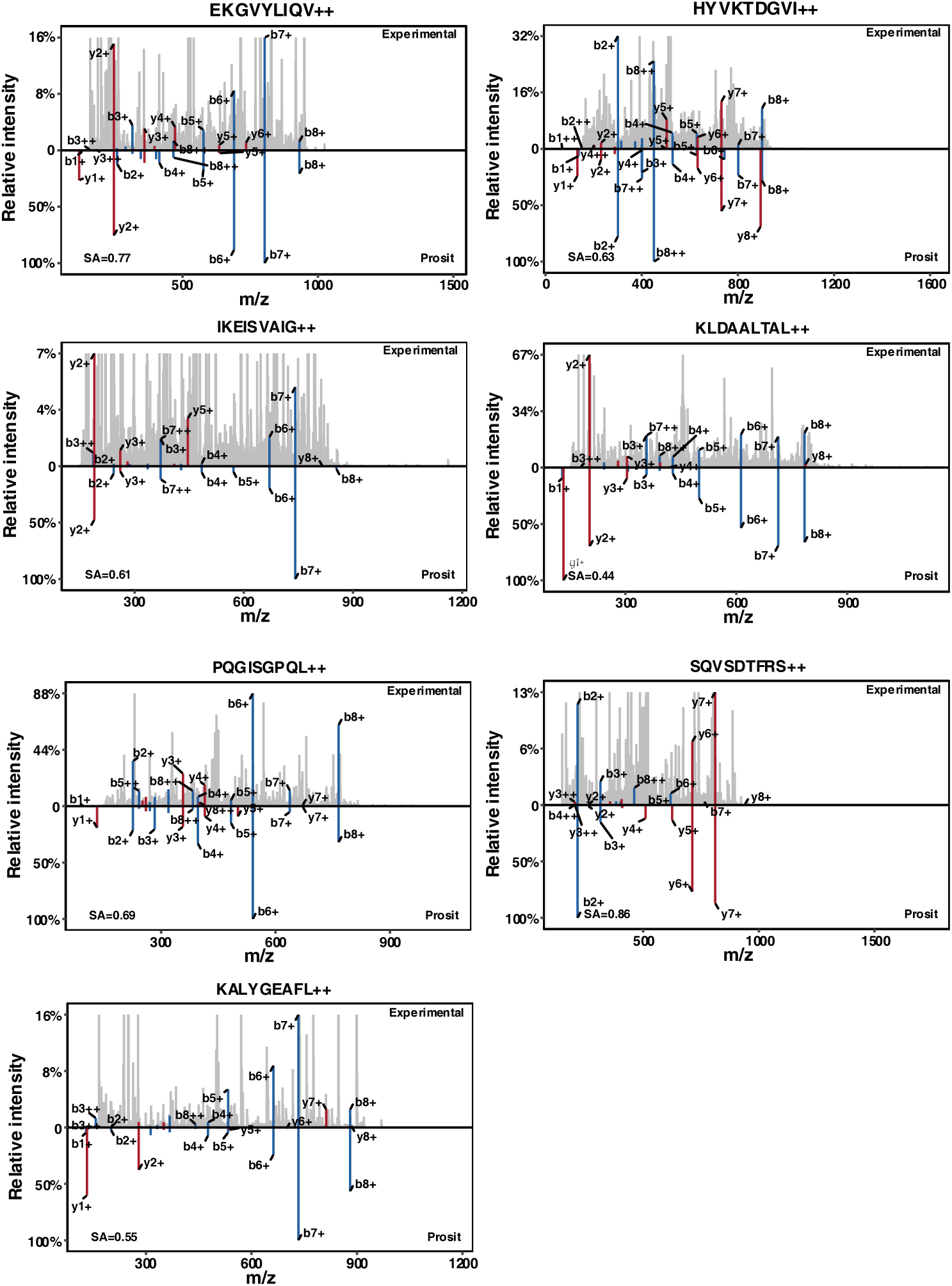
Mass spectra of the antigens. The top panel presents the experimental spectra acquired using DIA, while the bottom panel displays the predicted spectra generated by Prosit.

**Extended Data Fig. S16.**
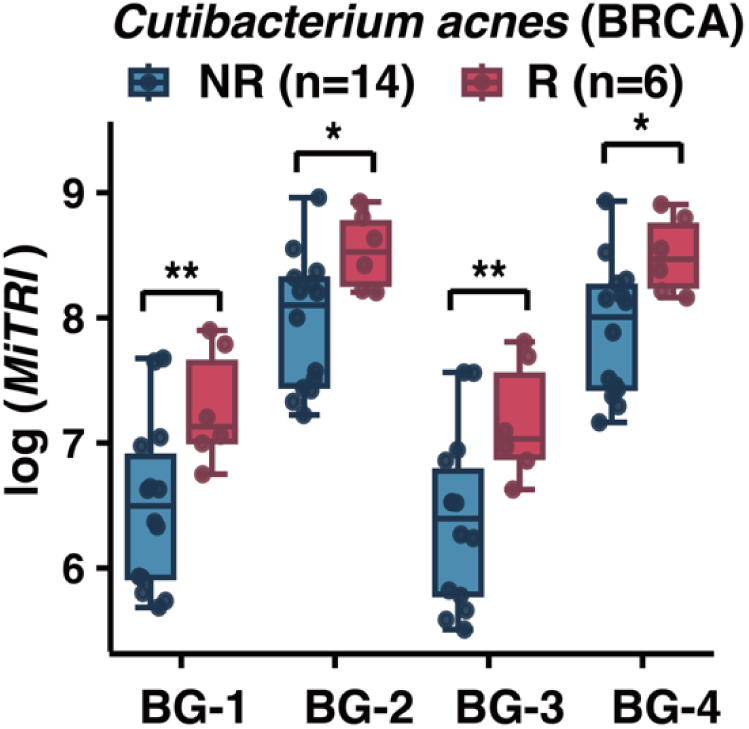
MiTRI of *Cutibacterium acnes* is significantly higher in neoadjuvant chemotherapy responders than non-responders in BRCA.

**Extended Data Fig. S17.**
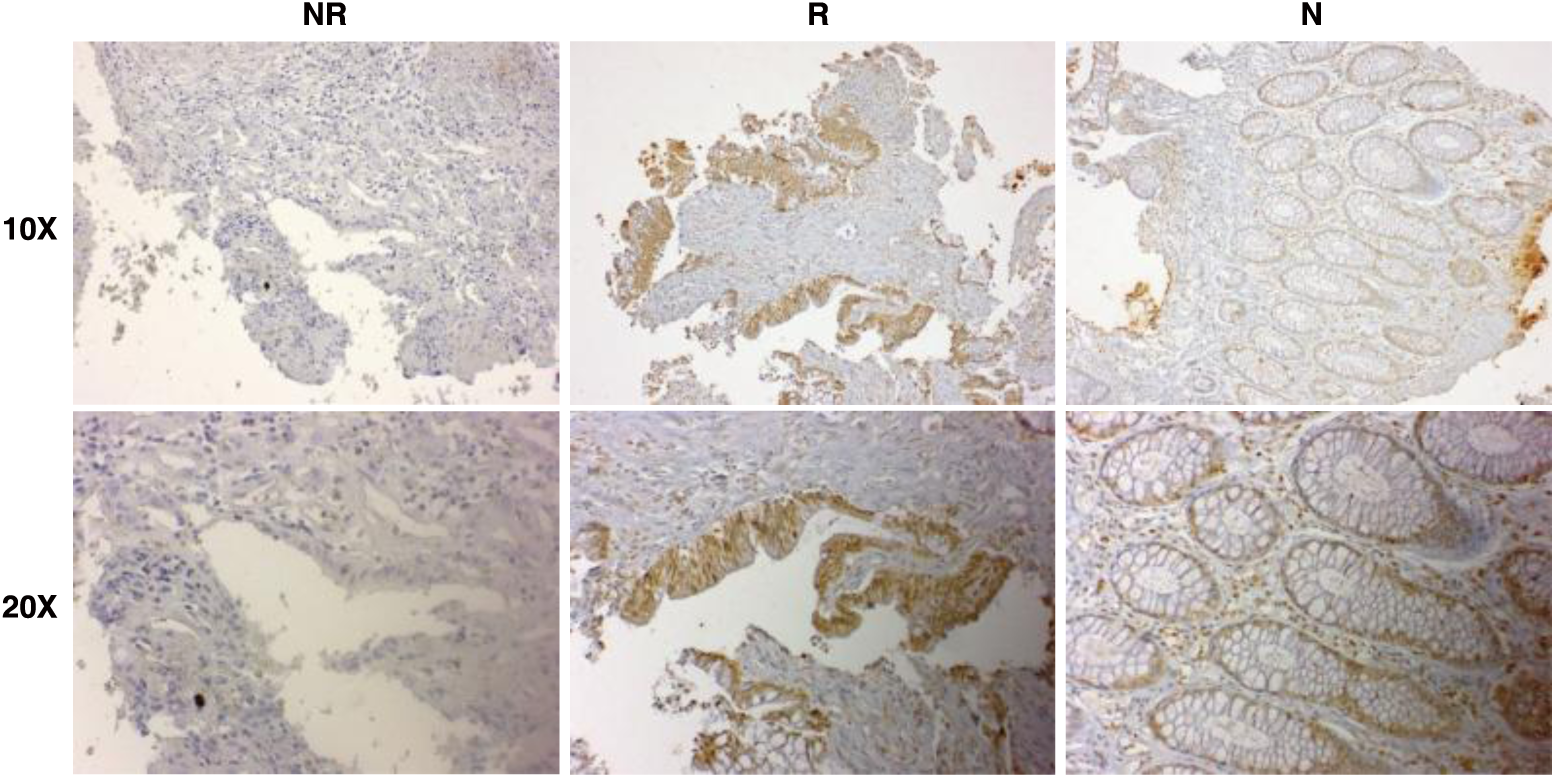
Immunohistochemistry staining for LPS reveals higher overall microbial abundance in neoadjuvant chemotherapy responders compared to non-responders in CRC.

**Extended Data Fig. S18.**
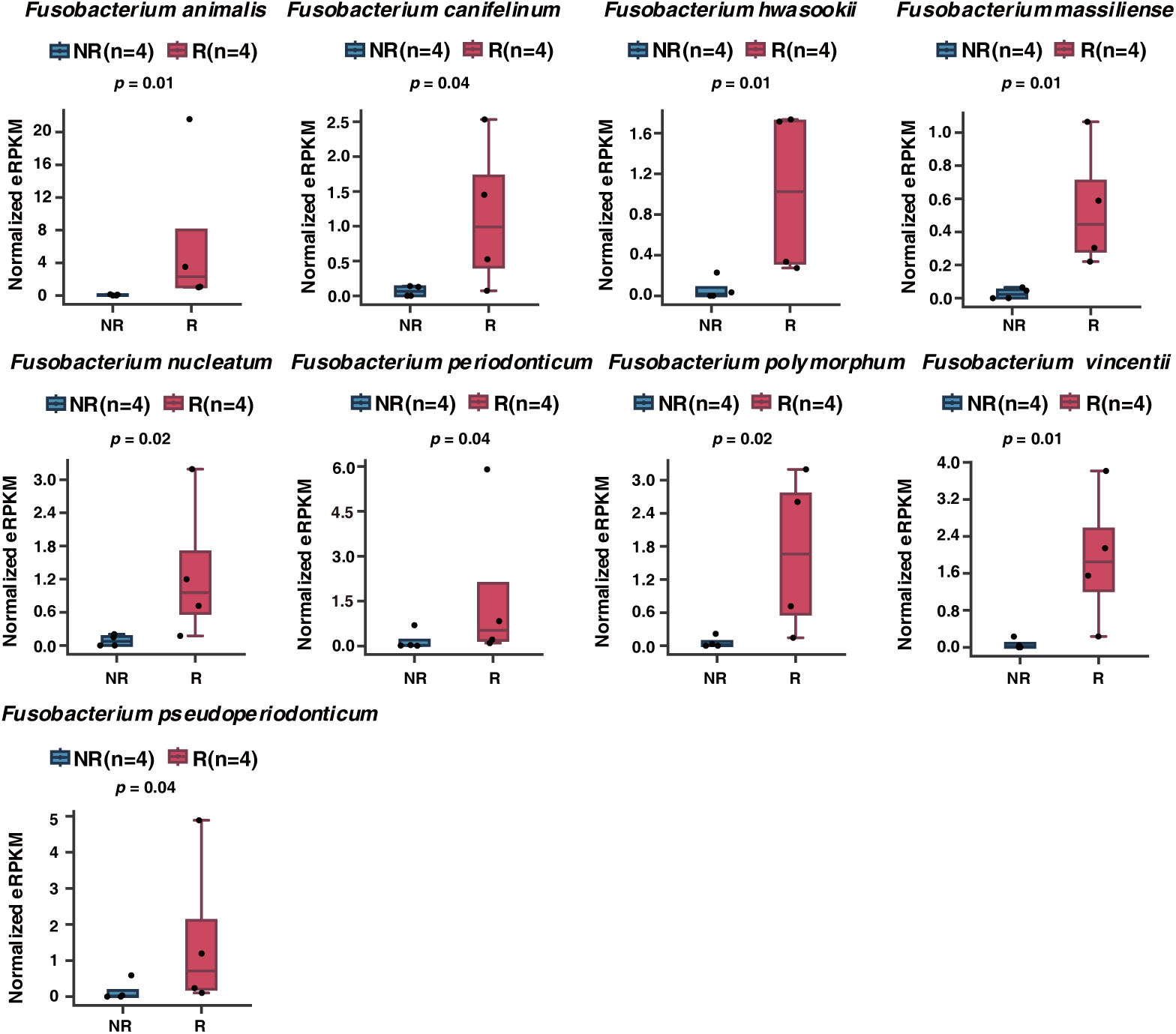
CRC-associated microbial abundance in neoadjuvant chemotherapy responders (R; n=4) and non-responders (NR; n=4) groups. Multiple CRC-associated microbial species exhibited significantly higher abundance in R group. P-values were calculated using one-sided Wilcoxon rank-sum ranksum tests.

**Extended Data Fig. S19.**
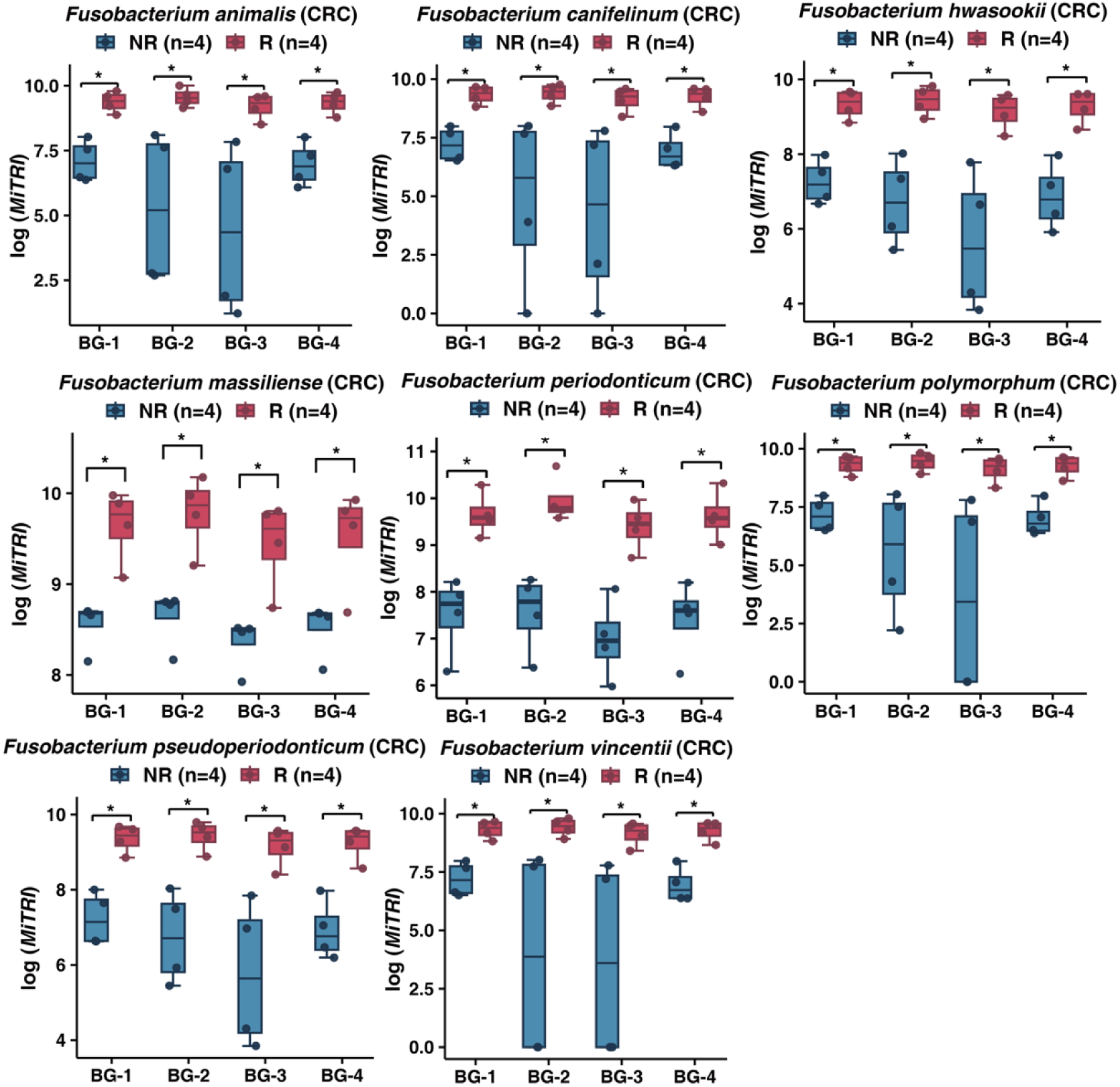
Microbial Transcriptional Reprogramming Index (MiTRI) of literature-reported CRC-associated microbes in neoadjuvant chemotherapy responders (R; n=4) and non-responders (NR; n=4) groups.

**Extended Data Fig. S20.**
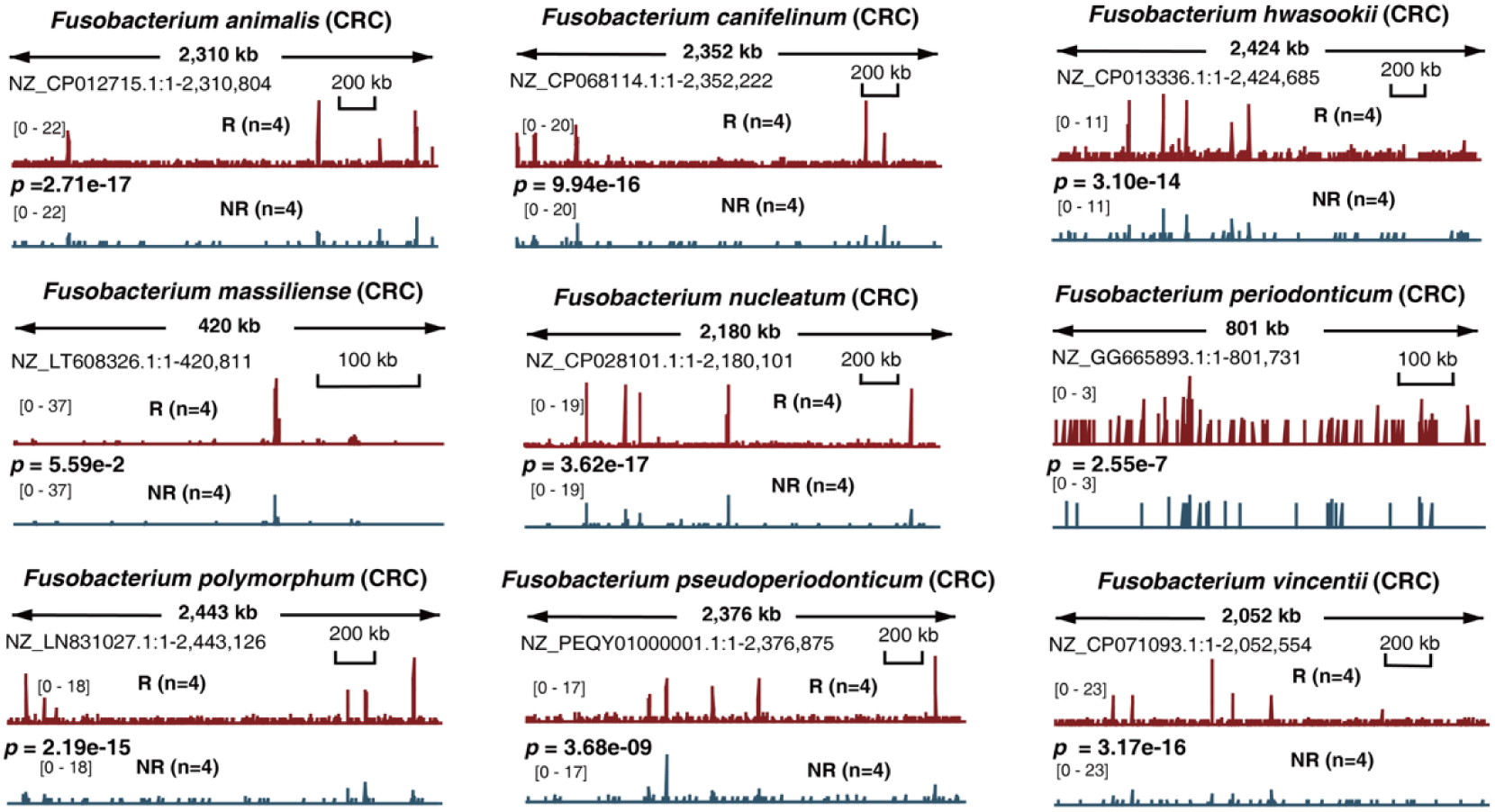
RNA-seq coverage analysis of CRC-associated microbes in neoadjuvant chemotherapy responders (R; n=4) and non-responders (NR; n=4) groups.

**Extended Data Fig. S21.**
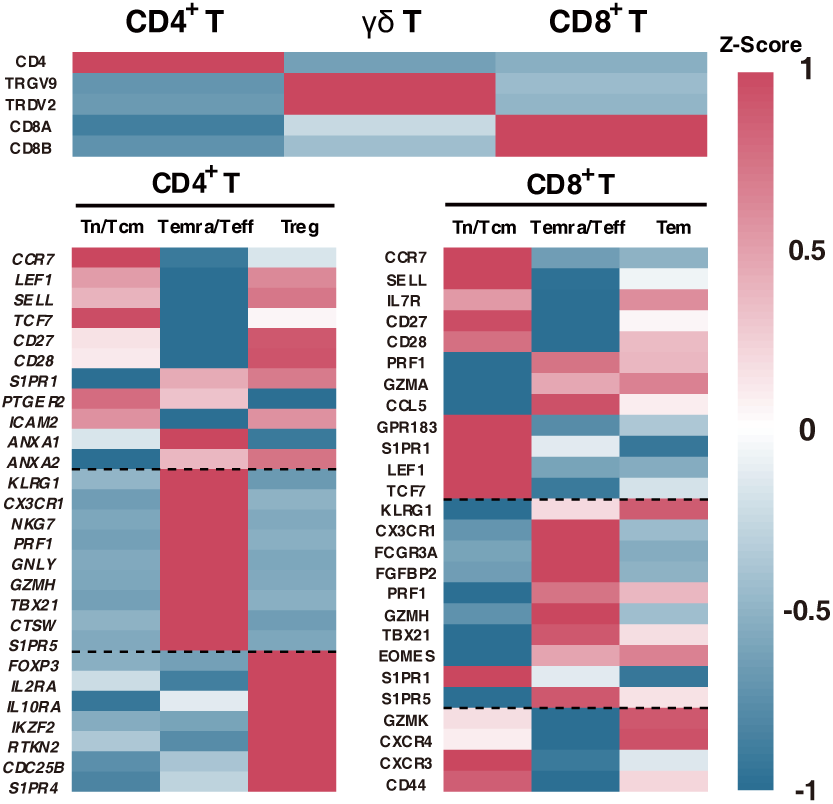
Heatmap of marker genes for single-cell annotation.

**Extended Data Fig. S22.**
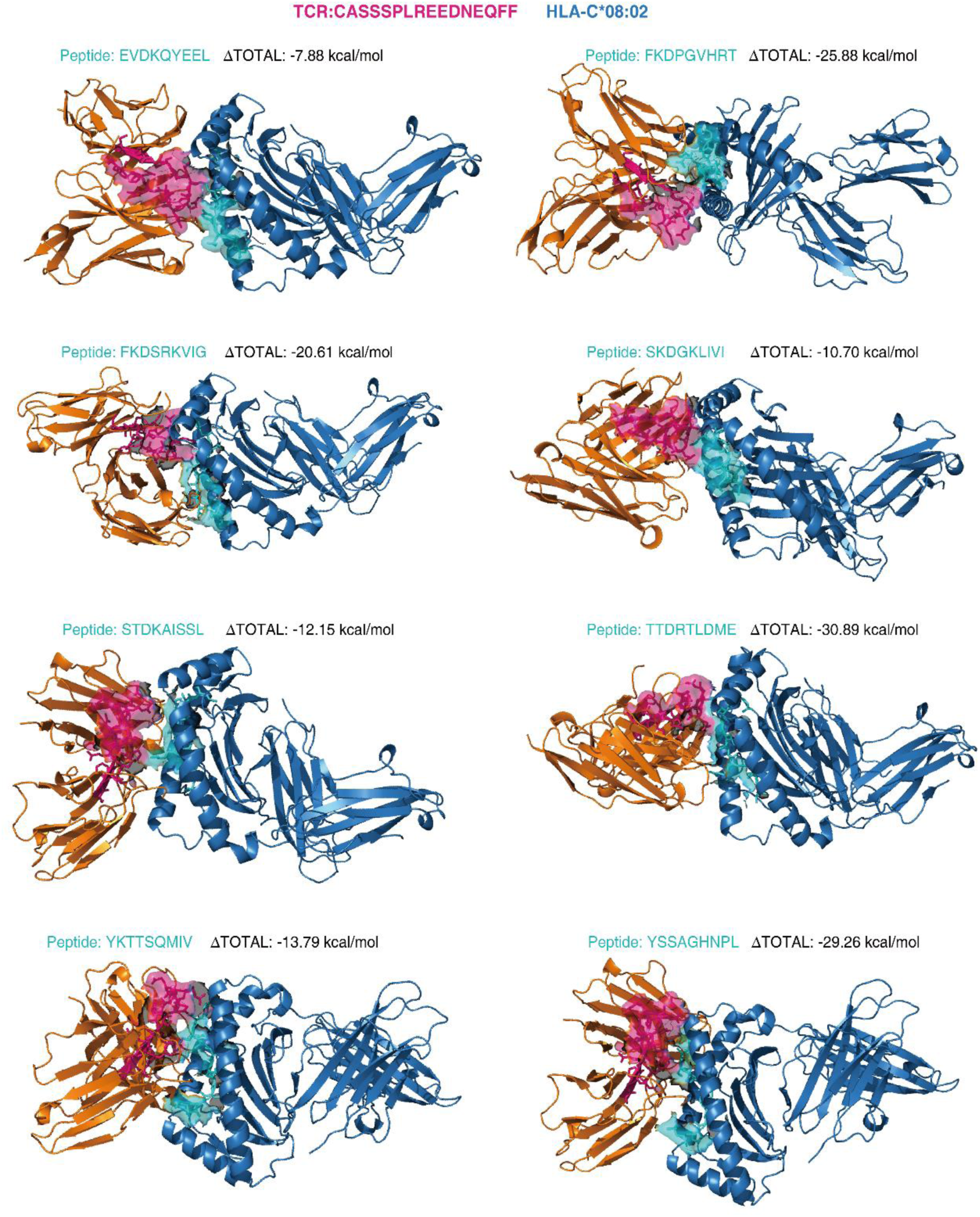
Structure-based binding energy analysis of eight peptides in complex with HLA-C*08:02 and the TCR CASSSPLREEDNEQFF. Binding energies were computed to evaluate the interaction strength and potential immunogenicity of each TCR–epitope–HLA complex.

**Extended Data Fig. S23.**
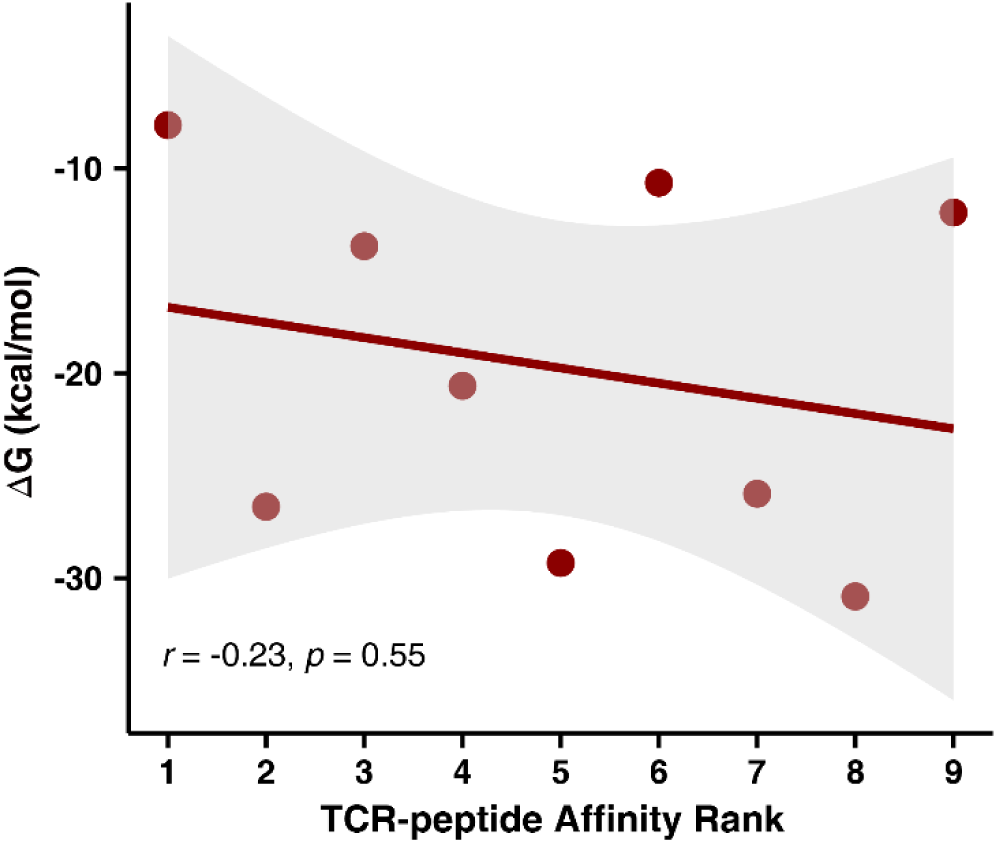
Correlation between the predicted TCR-binding affinity ranks and their binding energy (*r* = –0.23, *p* = 0.54, Pearson correlation).

**Extended Data Table S1. Metadata for publicly available and in-house sequencing datasets used in this study.** (Provided as a separate file)

**Extended Data Table S2. Custom dataset of RefSeq genomes derived from human-associated microbial catalogues.** (Provided as a separate file)

**Extended Data Table S3. Functional enrichment analysis of microbial expressed genes in peritumor and tumor tissues.** (Provided as a separate file)

**Extended Data Table S4. Immunopeptidomics validation of predicted microbial HLA-binding peptides.** (Provided as a separate file)

**Extended Data Table S5 List of literature-reported tumor-associated microorganisms in each cancer type** (Provided as a separate file)

**Extended Data Table S6. Colorectal cancer driver genes.** (Provided as a separate file)

**Extended Data Table S7. Single-cell annotation and analysis of T cell.** (Provided as a separate file)

**Extended Data Table S8. Predicted binding affinity between microbial candidate epitopes and TCR clonotypes.** (Provided as a separate file)

**Extended Data Table S9. Binding free energy between the TCR and epitope-HLA** (Provided as a separate file)

